# Environment-specific mechanosensing preserves a common bleb-based migratory program across diverse embryonic environments

**DOI:** 10.64898/2026.08.11.744326

**Authors:** Manami Morimoto, Yasuhiro Kamei, Mizuki Morita, Junichi Ikenouchi, Yoshiki Hayashi, Daisuke Saito

## Abstract

Cell migration frequently requires cells to traverse tissue environments with distinct physical and biochemical properties. How migrating cells preserve a common migratory program while adapting to these heterogeneous environments remains poorly understood. Here, we show that chick primordial germ cells (PGCs) preserve a common bleb-based migratory mode throughout embryogenesis despite migrating through mechanically distinct tissues. PGCs formed membrane blebs during both intravascular crawling and migration through the dorsal mesentery. However, nuclear envelope (NE) unfolding and activation of the NE-cPLA2 pathway occurred specifically during migration through the mechanically confined dorsal mesentery, where this pathway was required for bleb formation and efficient migration. In contrast, bleb formation during vascular crawling occurred independently of the NE-cPLA2 pathway, demonstrating that distinct molecular mechanisms can generate the same migratory behavior in different tissue environments. Together, these findings suggest that diverse environmental cues converge on a conserved bleb-forming machinery. We propose a hierarchical model in which migrating cells preserve a common migratory program by flexibly switching the upstream molecular mechanisms that initiate bleb formation according to the tissue environment.

## INTRODUCTION

Cell migration *in vivo* requires cells to traverse tissue environments that differ markedly in their physical and biochemical properties. During embryogenesis, immune surveillance, and cancer metastasis, migrating cells encounter diverse environments, including blood flow, endothelial barriers, extracellular matrices, and densely packed tissues (Bera et al., 2022; Friedl and Weigelin, 2008; Gensbittel et al., 2026; Huss et al., 2019; Yamada and Sixt, 2019). Despite these environmental transitions, many cell types migrate efficiently and reproducibly toward their destinations. How a single cell type preserves a robust migratory program while adapting to such heterogeneous environments remains a fundamental unresolved question.

Primordial germ cells (PGCs) provide an ideal model to address this question because they undergo long-distance migration during embryogenesis across a wide range of animal species before colonizing the developing gonads (Grimaldi and Raz, 2020). In avian embryos, this journey involves sequential migration through diverse environments, including the bloodstream, endothelial barriers, and densely packed mesenchymal tissues such as the dorsal mesentery (Morimoto and Saito, 2025; Morita et al., 2026; Saito et al., 2022). These environments impose distinct physical constraints, including blood flow, endothelial confinement, and tissue compression. Nevertheless, avian PGCs migrate with remarkable efficiency, suggesting that they possess mechanisms that preserve a common migratory behavior across diverse tissue environments.

Cell migration has traditionally been explained by chemotactic signaling pathways. In PGCs, guidance cues such as SDF-1/CXCR4, SCF/c-Kit, and non-canonical Wnt signaling have been implicated in migration (Doitsidou et al., 2002; Gu et al., 2009; Laird et al., 2011). It has become increasingly evident that mechanical cues also actively regulate cell migration through diverse mechanotransduction pathways. Cells sense mechanical stimuli through multiple mechanosensory systems, including mechanosensitive ion channels, integrin-mediated cell-matrix adhesions, and nuclear deformation (Canales Coutino and Mayor, 2021; Chastney et al., 2025; Kalukula et al., 2022; Penfield and Montell, 2023). Among these, nuclear envelope (NE) unfolding has emerged as a mechanosensory mechanism in which physical confinement stretches the NE, thereby activating cytosolic phospholipase A2 (cPLA2), and promoting actomyosin contractility and membrane bleb formation (Lomakin et al., 2020; Venturini et al., 2020). Although several mechanosensing pathways have been identified, whether migrating cells selectively employ distinct mechanosensory mechanisms according to their tissue environment remains largely unknown.

Membrane blebbing represents a highly efficient migration mode under mechanically confined conditions (Liu et al., 2015; Logue et al., 2015; Ruprecht et al., 2015). Bleb-based migration is driven by intracellular pressure and actomyosin contractility and has been observed in diverse biological contexts, including germ cell migration and cancer invasion (Blaser et al., 2006; Morita et al., 2025; Morita et al., 2026). Blebs are characterized by transient depletion of cortical actin and are frequently accompanied by localized Ca^2+^ signals (Aoki et al., 2021; Aoki et al., 2016; Charras and Paluch, 2008; Morita et al., 2026). Notably, we and others have shown that avian PGCs utilize bleb-based protrusions throughout multiple phases of their migratory journey (Ando and Fujimoto, 1983; Morita et al., 2026; Murai et al., 2021). Together, these observations raise the possibility that PGCs preserve a common bleb-based migratory mode across distinct tissue environments while engaging different mechanisms to initiate bleb formation.

Here, we show that avian PGCs preserve a common bleb-based migratory mode while employing distinct bleb-inducing mechanisms in different tissue environments. PGCs formed blebs during migration in both the vascular and mesenchymal phases, whereas NE unfolding occurred specifically during migration through the dorsal mesentery. Physical confinement selectively activated the NE-cPLA2 signaling pathway during migration through the dorsal mesentery, where it was required for bleb formation but dispensable within the vascular environment. Together, our findings support a hierarchical framework in which migrating cells preserve a common migratory program while flexibly engaging environment-specific mechanisms that initiate bleb formation.

## RESULTS

### PGCs preserve a common bleb-based migratory mode but exhibit context-dependent nuclear envelope unfolding

Our previous study demonstrated that chick PGCs utilize membrane blebs while crawling within the extravasation vascular plexus (Ex-VaP), a specialized capillary network where arrested PGCs initiate transendothelial migration into the dorsal mesentery (DM), a densely packed embryonic mesenchymal tissue (Morita et al., 2026). However, whether this bleb-based migratory mode is maintained after vascular exit into the mechanically distinct DM has remained unknown. To determine whether PGCs preserve a common bleb-based migratory mode across these two environments, we directly compared their behavior during the vascular and mesenchymal phases of migration.

Chicken PGCs can be maintained in long-term culture (Whyte et al., 2015), genetically manipulated *in vitro*, and subsequently transplanted into recipient embryos, allowing direct visualization of PGC behavior during migration *in vivo* (Saito et al., 2022). PGCs expressing the F-actin probe Lifeact-mCherry were transplanted into the bloodstream of Hamburger and Hamilton stage (HH)15 (Hamburger and Hamilton, 1951) chicken embryos, and their migratory behavior was analyzed by time-lapse imaging in both the Ex-VaP and the DM (Fig. 1a,b).

**Fig. 1.**
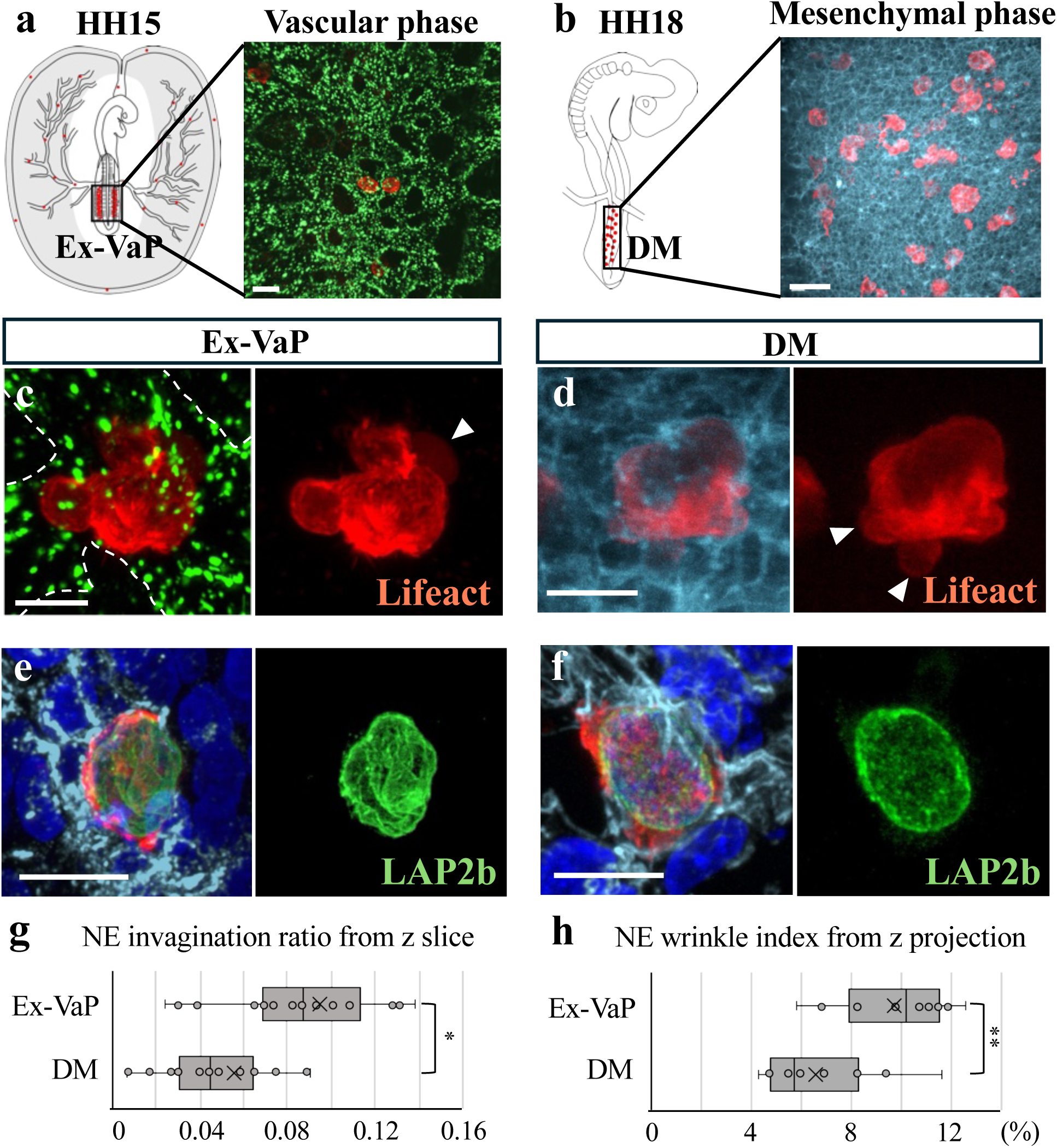
PGCs preserve a common bleb-based migratory mode while exhibiting context-dependent nuclear envelope unfolding. **a,b,** Schematic (left) and representative images (right) of the Ex-VaP at HH15 (a) and the DM at HH18 (b). **c,d,** Representative *ex vivo* live images of transplanted Lifeact-mCherry-expressing PGCs migrating within the Ex-VaP of an HH15 chick embryo (c) or the DM of an HH18 chick embryo (d). Endothelial cells were visualized by incorporation of 488-AcLDL (green) in c, and cell membranes were labeled with CellMask (cyan) in d. Arrowheads indicate membrane blebs lacking cortical F-actin. **e,f,** Representative confocal images of transplanted PGCs expressing LAP2b-AcGFP and Lifeact-mCherry within the Ex-VaP of an HH16 quail embryo (e) or the DM of an HH19 quail embryo (f). Endothelial cells were identified by QH1 immunostaining (cyan) in e, whereas F-actin was visualized with Phalloidin (cyan) in f. **g, h,** Quantification of the NE invagination ratio (g) and NE wrinkling index (h) in PGCs within the Ex-VaP and the DM (Ex-VaP: n = 22 cells from 6 embryos; DM: n = 27 cells from 4 embryos). Scale bars, 30 µm (a, b); 10µm (c-f). *P<0.05, **P<0.01 (two-sided unpaired Student’s t-test).

Consistent with our previous observation, PGCs crawling within the Ex-VaP formed balloon-like protrusions lacking cortical F-actin, characteristic of membrane blebs (Fig. 1c). Remarkably, PGCs migrating through the DM likewise generated blebs throughout migration (Fig. 1d). Similar bleb structures were also observed in endogenous PGCs within both the Ex-VaP and the DM (Extended Data Fig. 1a,b). Together, these observations indicate that chick PGCs utilize blebs in both vascular and mesenchymal environments despite their markedly different physical properties.

Recent studies have shown that NE unfolding promotes bleb formation in physically confined cells (Lomakin et al., 2020; Venturini et al., 2020). We therefore asked whether this mechanosensing mechanism is similarly engaged in both environments. To determine whether NE morphology differs between them, PGCs expressing AcGFP-tagged LAP2b, an inner nuclear membrane protein, were transplanted into quail embryos, in which the vascular endothelium can be specifically identified by QH1 immunostaining (Pardanaud et al., 1987). This approach allowed accurate discrimination between PGCs within the Ex-VaP and those migrating through the DM. Because NE unfolding is reflected by both large-scale changes in nuclear shape and fine-scale alterations in NE surface topology, we quantified NE morphology using two complementary metrics. The NE invagination ratio measures large inward deformations of the NE from z-sections (Venturini et al., 2020), whereas the wrinkling index quantifies fine-scale NE folding from z-projection images (Cosgrove et al., 2021).

PGCs arrested within the Ex-VaP retained highly folded nuclear envelopes (Fig. 1e), whereas PGCs migrating through the DM displayed markedly unfolded nuclear envelopes (Fig. 1f). Consistent with these observations, both the NE invagination ratio and the wrinkling index were significantly reduced in DM PGCs compared with those arrested within the Ex-VaP (Fig. 1g,h). Together, these findings demonstrate that chick PGCs preserve a common bleb-based migratory program while selectively undergoing NE unfolding during migration through the physically confined DM.

### *In vitro* confinement recapitulates the mechanical environment of PGCs in the dorsal mesentery

The tissue-specific NE unfolding observed during migration through the DM suggested that PGCs experience a distinct mechanical environment in this tissue. We therefore asked whether this environment could be recapitulated *in vitro* using our previously established under-agarose confinement assay (Morita et al., 2026), in which cultured PGCs are mechanically compressed between a glass coverslip and an agarose gel pad. Confinement height was controlled using microbeads of defined diameters (Extended Data Fig. 2a,b).

We first asked whether physical confinement was sufficient to reproduce the NE morphology observed *in vivo*. Confinement at both 10-μm and 7-μm height induced membrane blebbing, and the proportion of blebbing cells increased as confinement height was reduced (Fig. 2a,b). Likewise, reducing the confinement height progressively promoted NE unfolding, as reflected by decreases in the NE invagination ratio and wrinkling index (Fig. 2c,d). Notably, the extent of NE unfolding under the 10-μm confinement condition closely resembled that observed in PGCs migrating through the DM, whereas the 7-μm condition produced more extensive NE unfolding. These findings identify the 10-μm confinement condition as the *in vitro* model that most closely reproduces the nuclear morphology of PGCs in the DM.

**Fig. 2.**
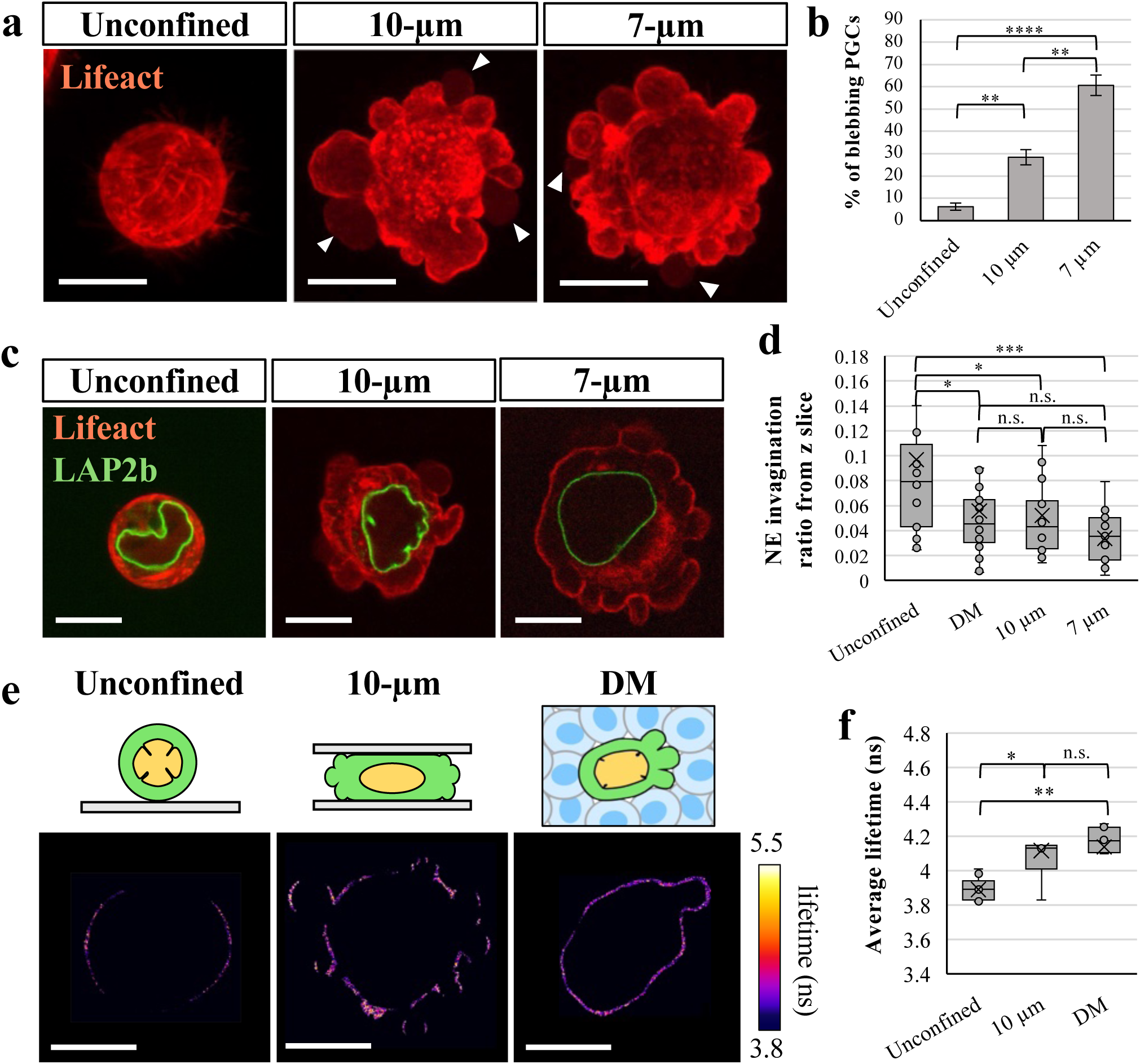
*In vitro* confinement recapitulates the mechanical environment of PGCs in the DM. **a,b,** Representative images (a) and quantification (b) of Lifeact-mCherry-expressing PGCs under unconfined conditions or confined beneath an agarose gel at 10-µm, or 7-µm heights. Arrowheads indicate membrane blebs lacking cortical F-actin. Data are presented as mean ± SD from three independent experiments (n = 3), with approximately 50 cells analyzed per experiment. **c,d,** Representative z-section images (c) and quantification (d) of NE morphology in PGCs expressing Lifeact-mCherry and LAP2b-AcGFP under unconfined conditions, in the DM, or under 10-µm or 7-µm confinement (Unconfined: n = 21 cells; DM: n = 27 cells; 10-µm: n = 20 cells; 7-µm: n = 19 cells). **e,f,** Representative FLIM images (e) and quantification (f) of Flipper-TR fluorescence lifetime measured at the plasma membranes of PGCs under unconfined conditions, under 10-µm confinement, or during migration in the DM. Quantification was performed from three independent experiments (Unconfined: n = 9 cells; 10-µm: m = 7 cells; DM: n = 8 cells). Scale bars, 10 µm. n.s., not significant; *P<0.05, **P<0.01 (two-sided unpaired Student’s t-test).

We next asked whether the 10-μm confinement condition also recapitulates the mechanical state experienced by PGCs *in vivo*. As membrane tension is a key physical parameter reflecting cellular mechanical state, we quantified membrane tension using Flipper-TR, a mechanosensitive fluorescent probe whose fluorescence lifetime increases with membrane tension (Colom et al., 2018; Luchtefeld et al., 2024; Wang et al., 2024). Fluorescence lifetime imaging microscopy (FLIM) revealed that membrane tension was significantly increased under the 10-μm confinement condition compared with unconfined controls (Fig. 2e,f). Importantly, the fluorescence lifetime of confined PGCs closely matched that of PGCs migrating through the DM, indicating that the 10-μm confinement condition faithfully reproduces the mechanical state of the DM.

Together, these findings demonstrate that the 10-μm confinement assay recapitulates both the nuclear morphology and mechanical state of PGCs migrating through the DM. This physiologically relevant system therefore provides a robust platform for dissecting the molecular mechanisms by which physical confinement promotes bleb formation.

### Physical confinement selectively activates the NE-cPLA2 pathway, whereas cortical myosin II recruitment accompanies bleb formation in both migratory environments

Having established a physiologically relevant confinement assay that recapitulates the mechanical environment of the DM, we next asked whether physical confinement activates the NE-cPLA2 signaling pathway previously described in mechanically confined cells. In this pathway, mechanical unfolding of the NE activates cPLA2, which in turn promotes cortical actomyosin assembly and membrane bleb formation (Lomakin et al., 2020; Venturini et al., 2020).

To determine whether this pathway is activated in PGCs, cultured cells were subjected to the 10-μm confinement assay established in Figure 2. Consistent with previous reports, physical confinement induced prominent nuclear accumulation of cPLA2 (Fig. 3a-c) (Alraies et al., 2024). In parallel, myosin II became enriched at the cell cortex, and the cortical intensity of its cortical intensity increased significantly under confined conditions (Fig. 3d-f). These observations demonstrate that the physiologically relevant confinement condition activates the canonical NE-cPLA2 signaling pathway in cultured PGCs.

**Fig. 3.**
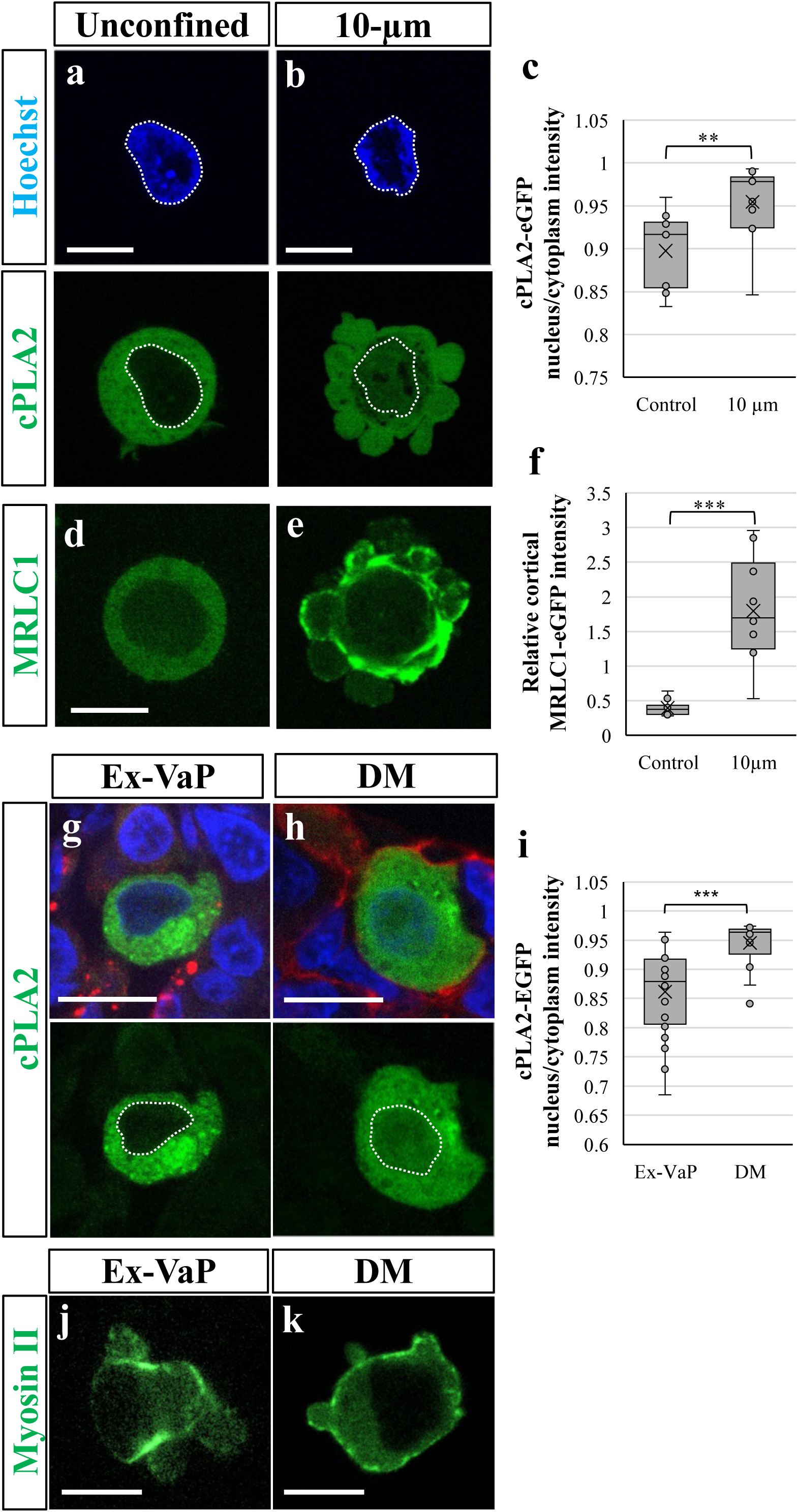
Physical confinement activates the NE-cPLA2 signaling pathway *in vitro*. **a-c,** Representative z-section images (a,b) and quantification of the nuclear-to-cytoplasmic ratio of cPLA2-EGFP fluorescence intensity (c) in PGCs under unconfined conditions (a) or under 10-µm confinement (b). Nuclei were stained with Hoechst. Dotted lines outline the nucleus (Unconfined: n = 10 cells; 10-µm: n = 13 cells). **d-f,** Representative z-section images (d,e) and quantification of relative MRLC1-EGFP enrichment (f) under unconfined conditions (d) or under 10-µm confinement (e) (n = 10 cells for each condition). **g-i,** Representative transverse sections (g,h) and quantification of nuclear-to-cytoplasmic ratio of cPLA2-EGFP fluorescence intensity (i) in transplanted PGCs within the Ex-VaP of HH16 chick embryos (g) or the DM of HH19 chick embryos (h). Endothelial cells were visualized by incorporation of 594-AcLDL (red) in g, whereas F-actin was visualized with phalloidin (red) in h. Nuclei were stained with DAPI (blue). Dotted lines outline the nucleus (Ex-VaP: n = 16 cells from 5 embryos; DM: n = 17 cells from 5 embryos). **j**, **k**, Representative z-section images of transplanted PGCs expressing MRLCI-EGFP (Myosin II) within the Ex-VaP (j) or the DM (k). Scale bars, 10 µm. **P<0.01, ***P<0.001 (two-sided unpaired Student’s t-test).

We next asked whether these responses also occur during PGC migration *in vivo*. PGCs expressing fluorescently labeled cPLA2 or myosin II were transplanted into recipient embryos, and their subcellular localization was examined in the Ex-VaP and the DM (Fig. 3g-k). Whereas cPLA2 was predominantly cytoplasmic in PGCs arrested within the Ex-VaP, PGCs migrating through the DM exhibited marked nuclear accumulation of cPLA2 (Fig. 3g-i). In contrast, cortical accumulation of myosin II was observed in both migratory environments (Fig. 3j,k), consistent with the occurrence of membrane blebbing in both the Ex-VaP and the DM. These localization patterns indicate that whereas cortical actomyosin recruitment is a shared feature of bleb-based migration, activation of the NE-cPLA2 pathway is selectively associated with migration through the mechanically confined DM.

Together, these findings demonstrate that physical confinement selectively activates the NE-cPLA2 signaling pathway both *in vitro* and *in vivo*, while cortical recruitment of myosin II accompanies bleb formation in both vascular and mesenchymal environments. The close correspondence between the *in vitro* confinement model and PGCs migrating through the DM further establishes this assay as a physiologically relevant platform for dissecting the molecular mechanisms underlying confinement-induced bleb formation.

### The NE-cPLA2 signaling pathway is required for confinement-induced bleb formation

Having shown that physical confinement activates the NE-cPLA2 signaling pathway, we next asked whether this pathway is required for confinement-induced bleb formation.

To inhibit NE unfolding, we overexpressed lamin B receptor (LBR), an inner nuclear membrane protein that promotes NE folding by strengthening the association between the nuclear membrane and chromatin (Ma et al., 2007). LBR expression was induced using a Tet-on inducible expression system (Watanabe et al., 2007) by treating cells with doxycycline (Dox) for 3 days before analysis. As expected, LBR-overexpressing (LBR-OE) PGCs exhibited highly folded nuclear envelopes under unconfined conditions (Fig. 4a-c). Under 10-μm confinement, control PGCs underwent marked NE unfolding, whereas LBR-OE PGCs retained highly folded nuclear envelopes (Fig. 4d-f). Importantly, LBR overexpression significantly suppressed confinement-induced bleb formation (Fig. 4g,h). These findings demonstrate that NE unfolding is required for bleb formation under physiologically relevant confinement.

**Fig. 4.**
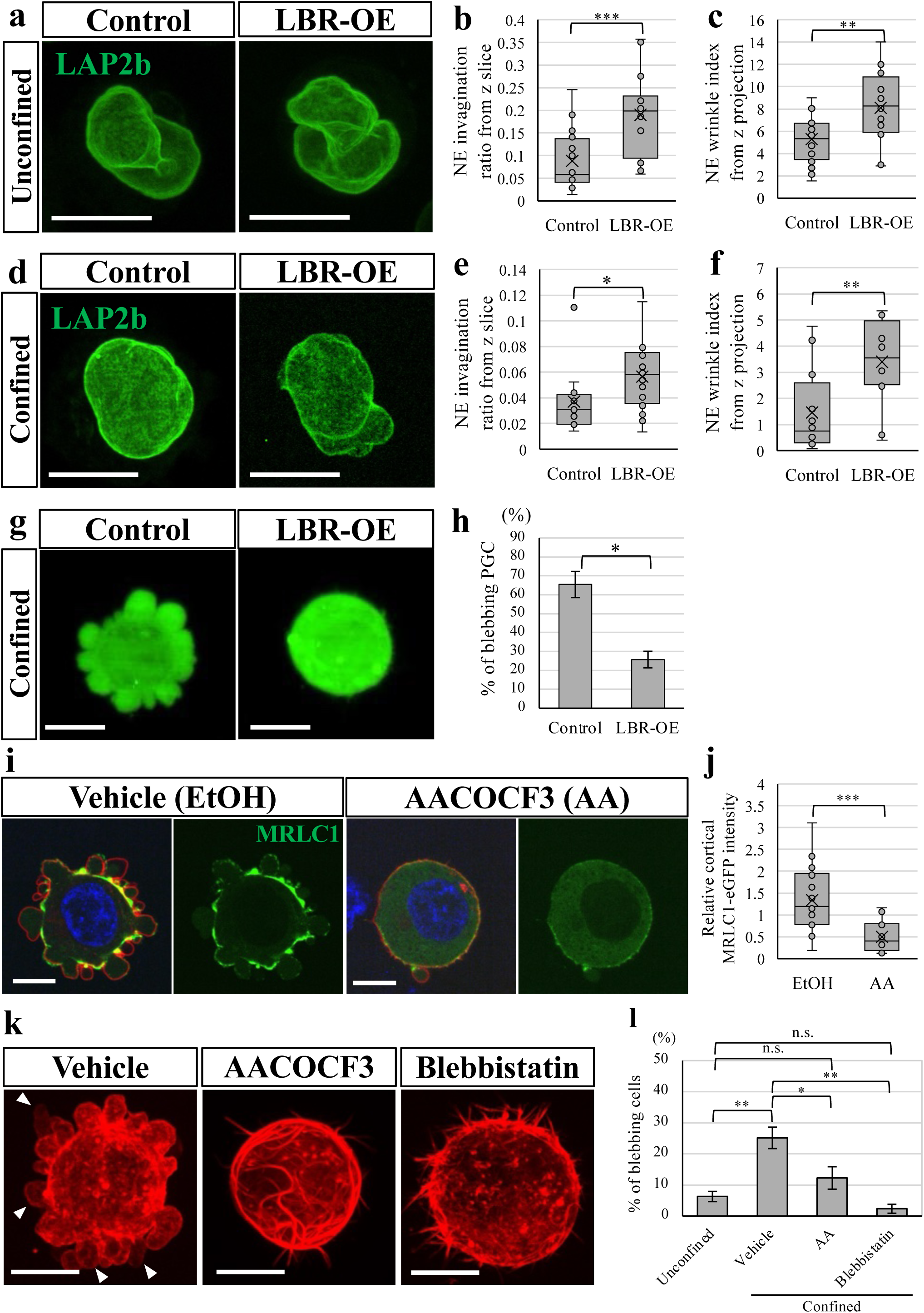
The NE-cPLA2 signaling pathway is required for confinement-induced bleb formation *in vitro*. **a-c,** Representative z-section images (a) and quantification of the NE invagination ratio (b) and NE wrinkling index (c) in control and LBR-OE PGCs under unconfined conditions (Control: n = 19 cells; LBR-OE: n = 20 cells). **d-f,** Representative z-section images (d) and quantification of the NE invagination ratio (e) and NE wrinkling index (f) in control and LBR-OE PGCs under 10-µm confinement (Control: n =12 cells; LBR-OE: n = 20 cells). **g,h,** Representative z-section images (g) and quantification (h) of bleb formation in control (ZsGreen1) and LBR-OE (ZsGreen1+LBR) PGCs under 10-µm confinement. Data are presented as mean ± SD from three independent experiments (n = 3), approximately 30 cells analyzed per experiment. **i,j,** Representative z-section images (i) and quantification (j) of control accumulation of MRLCI-EGFP (Myosin II) in PGCs under 10-µm confinement treated with vehicle (EtOH) or the cPLA2 inhibitor AACOCF3. Cell membranes were visualized with mCherry-CAAX (red), and nuclei were stained with DAPI (blue) (n = 18 cells for each condition). **k,l,** Representative images (k) and quantification (l) of bleb formation in Lifeact-mCherry-expressing PGCs under 10-µm confinement treated with vehicle (EtOH), AACOCF3, or blebbistatin. Arrowheads indicate membrane blebs lacking cortical F-actin. Data are presented as the mean ± SD from three independent experiments (N = 3), with approximately 30 cells analyzed per experiment. Scale bars, 10 µm. *P<0.05, **P<0.01, ***P<0.001 (two-sided unpaired Student’s t-test).

We next examined the requirement for cPLA2, a key effector of the NE signaling pathway. Pharmacological inhibition of cPLA2 with arachidonyl trifluoromethyl ketone (AACOCF3; AA) markedly reduced cortical accumulation of myosin II and significantly suppressed bleb formation under confined conditions (Fig. 4i-l). These findings indicate that cPLA2 couples NE-dependent mechanosensing to cortical actomyosin assembly.

Finally, we asked whether cortical actomyosin contractility is required for confinement-induced bleb formation. Pharmacological inhibition of myosin II with blebbistatin markedly reduced the proportion of blebbing PGCs under confined conditions (Fig. 4k,l), demonstrating that cortical actomyosin contractility is essential for bleb formation induced by physiological confinement. Together, these findings identify place cPLA2 downstream of NE mechanosensing and upstream of cortical actomyosin assembly.

### The NE-cPLA2 signaling pathway is specifically required for bleb formation during migration through the dorsal mesentery

Having established that the NE-cPLA2 signaling pathway is required for confinement-induced bleb formation *in vitro*, we next asked whether this pathway is similarly required during PGC migration through distinct tissue environments *in vivo*.

To determine the requirement for NE unfolding, we overexpressed LBR, which maintains a highly folded nuclear envelope under confinement, and analyzed PGC behavior following transplantation into recipient embryos. Time-lapse imaging revealed that approximately half of control PGCs formed blebs while migrating through the DM, whereas LBR-OE PGCs exhibited a marked reduction in bleb formation (Fig. 5a,b). In striking contrast, LBR overexpression had no detectable effect on bleb formation by PGCs crawling within the Ex-VaP (Fig. 5c,d). These findings demonstrate that NE unfolding is specifically required for bleb formation during migration through the mechanically confined DM but is dispensable during vascular crawling.

**Fig. 5.**
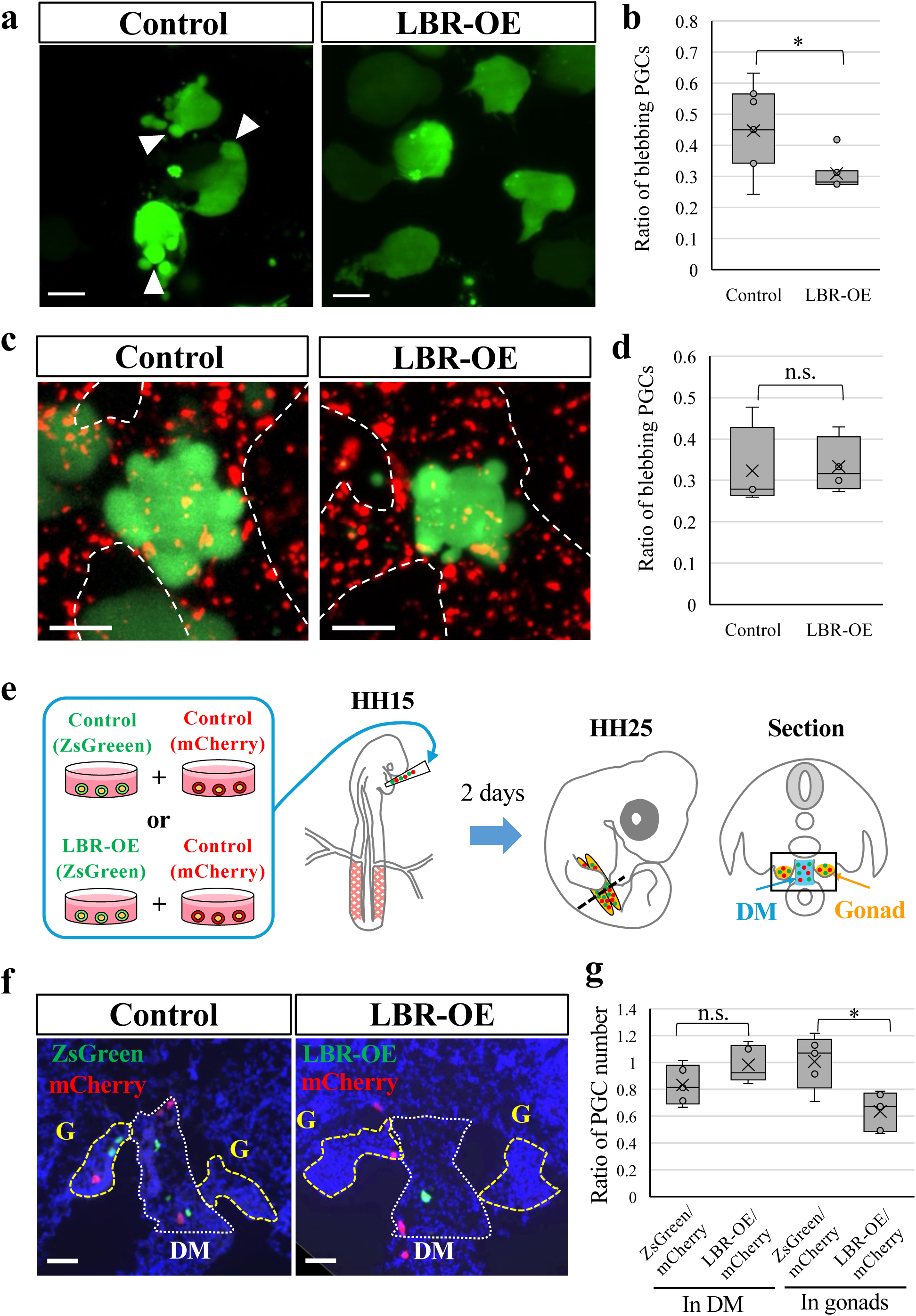
NE unfolding is specifically required for bleb formation and efficient migration through the dorsal mesentery. **a,b,** Representative *ex vivo* live images (a) and quantification (b) of bleb formation in control (ZsGreen1) and LBR-OE (LBR-OR; ZsGreen1+LBR) PGCs migrating through the DM of HH18 chick embryos. Arrowheads indicate membrane blebs (n = 7 embryos per condition; at least 20 cells were analyzed per embryo). **c,d,** Representative *ex vivo* live images (c) and quantification (d) of bleb formation in control and LBR-OE PGCs crawling within the Ex-VaP of HH15 chick embryos. Endothelial cells were visualized by incorporation of 594-AcLDL (red). Dotted lines outline endothelial cells. (n = 4 embryos per condition; at least 20 cells were analyzed per embryo). **e**, Schematic of the experimental design for analysis of PGC migration *in vivo*. An equal number of control (ZsGreen1) or LBR-OE (ZsGreen1+LBR) PGCs were co-transplanted with control mCherry-labeled PGCs into HH15 chick embryos. At HH25, the number of ZsGreen1-positive and mCherry-positive PGCs was quantified in the DM and gonads. **f,g,** Representative transverse sections (f) and quantification (g) of transplanted PGC distribution at HH25. Yellow and white dotted lines indicate the gonads (G) and DM, respectively. Data were obtained from n = 5 embryos, with at least 500 cells analyzed per embryo. Scale bars, 10 µm (a, c); 50µm (f). n.s., not significant. *P<0.05, **P<0.01, ***P<0.001 (two-sided paired Student’s t-test).

To determine whether this environment-specific requirement contributes to migratory success, equal numbers of control and LBR-OE PGCs were co-transplanted into HH15 chicken embryos, and their distribution was analyzed at HH25 (Fig. 5e). Significantly fewer LBR-OE PGCs reached the developing gonads, whereas a greater proportion remained within the DM (Fig. 5f,g). These findings indicate that NE unfolding is specifically required for efficient migration through the DM toward the gonads. Importantly, LBR overexpression did not affect PGC proliferation during culture or arrest within the Ex-VaP, indicating that the migratory defect was not secondary to impaired cell growth or defective vascular arrest (Extended Data Fig. 3a-c).

We next asked whether cPLA2 exhibits the same environment-specific requirement. Complete disruption of cPLA2 by CRISPR-Cas9-mediated genome editing resulted in rapid cell death during culture, precluding subsequent migration analyses. We therefore performed cPLA2 knockdown by an RNAi system modified for chicken (Das et al., 2006). However, knockdown efficiency was limited (∼50%), and cPLA2 knockdown alone did not significantly suppress confinement-induced bleb formation (Extended Data Fig. 4a-c). To overcome this limitation, we combined RNAi-mediated partial knockdown with transient treatment using the irreversible cPLA2 inhibitor AA to suppress cPLA2 activity during the migration period. The suppression of confinement-induced bleb formation persisted for at least 12 h after drug removal, enabling subsequent transplantation analyses (Extended Data Fig. 4d,e).

Using this approach, we found that cPLA2 inhibition phenocopied LBR overexpression. Bleb formation was significantly reduced during migration through the DM (Fig. 6a,b) but was unaffected during vascular crawling within Ex-VaP (Fig. 6c,d). Likewise, the analysis of PGC distribution at HH21 revealed that cPLA2-inhibited PGCs exhibited impaired migration toward the developing gonads following vascular exit (Extended Data Fig. 5a and Fig. 6e,f). Importantly, cell proliferation remained unaffected for at least one day after drug removal, indicating that the migratory defect was not secondary to impaired cell growth (Extended Data Fig. 5b).

**Fig. 6.**
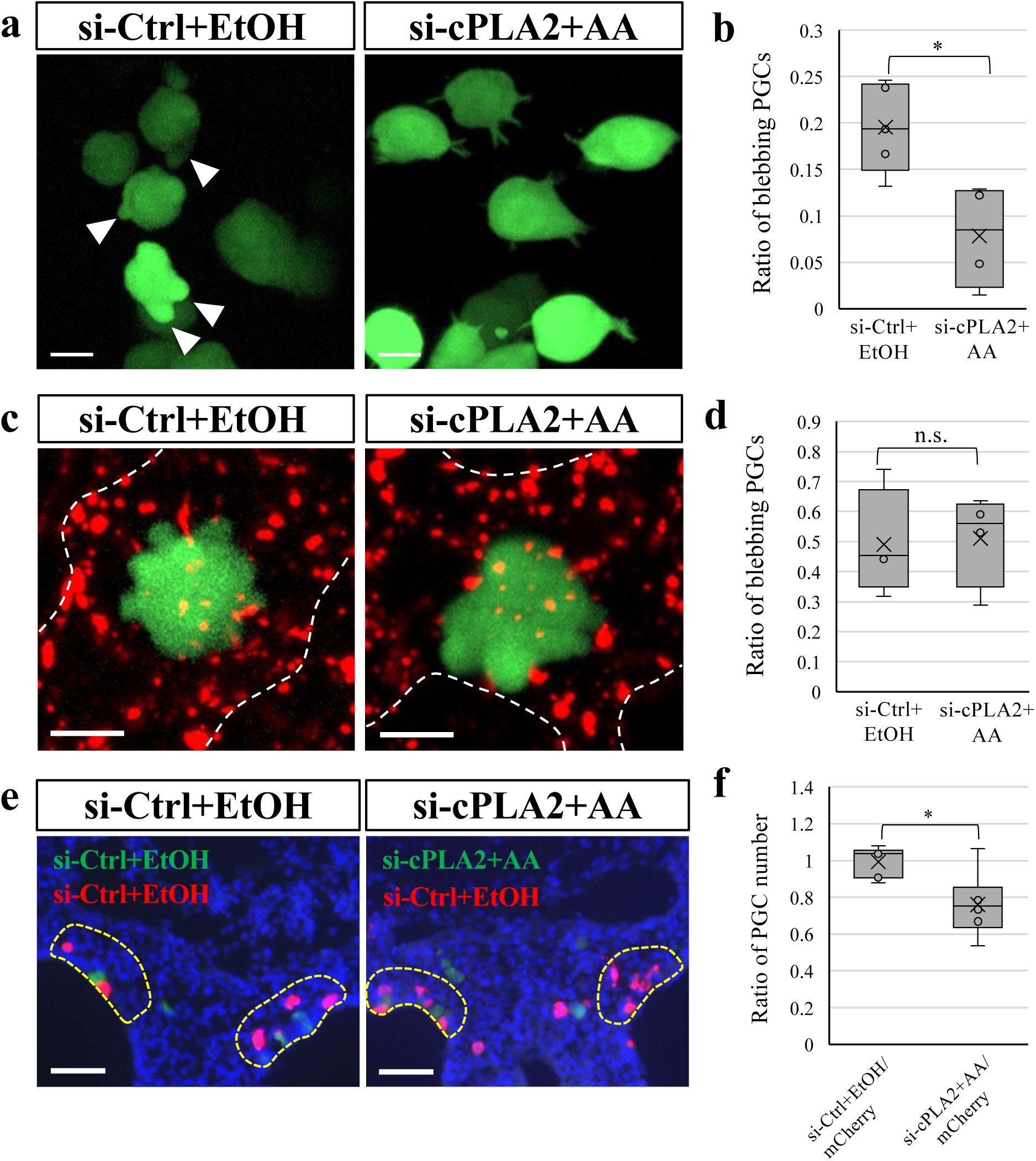
cPLA2 activity is specifically required for bleb formation and efficient migration through the dorsal mesentery. **a,b,** Representative *ex vivo* live images (a) and quantification (b) of bleb formation in si-Ctrl +EtOH and si-cPLA2+AACOCF3 (AA) PGCs migrating through the DM of HH18 chick embryos. Arrowheads indicate membrane blebs (si-Ctrl+EtOH: n = 5 embryos; si-cPLA2+AA: n = 4 embryos; at least 30 cells were analyzed per embryo). **c,d,** *ex vivo* live images (c) and quantification (d) of bleb formation in si-Ctrl+EtOH and si-cPLA2+AA PGCs crawling within the Ex-VaP in HH15 embryos. Endothelial cells were visualized by incorporation of 594-AcLDL (red). Dotted lines outline endothelial cells (n = 4 embryos per condition; at least 20 cells were analyzed per embryo). **e,f,** Representative transverse sections (e) and quantification (f) of transplanted PGC distribution in the gonads of HH21 chick embryos. Control (si-Ctrl+EtOH; EGFP) or cPLA2-inhibited (si-cPLA2+AA; EGFP) PGCs were co-transplanted with control mCherry-labeled PGCs at HH15. Yellow dotted lines indicate the gonads. Data were obtained from si-Ctrl+EtOH/mCherry (n = 7 embryos) and si-cPLA2+AA/mCherry (n = 6 embryos), with at least 500 PGCs analyzed per embryo. Scale bars, 10 µm (a, c); 50µm (e). n.s., not significant. *P<0.05, **P<0.01, ***P<0.001 (two-sided unpaired Student’s t-test).

Finally, PGCs continued to form blebs while crawling within the Ex-VaP following experimental cessation of blood flow (Extended Data Fig. 6a-d), indicating that continuous blood flow is not required for vascular bleb formation. Together with the insensitivity of vascular blebbing to both LBR overexpression and cPLA2 inhibition, these findings indicate that bleb formation within the Ex-VaP is regulated independently of the confinement-induced NE-cPLA2 pathway.

Together, these findings demonstrate that the NE-cPLA2 pathway is specifically required for bleb formation during migration through the mechanically confined DM, whereas vascular blebbing is likely regulated by a distinct mechanism independent of confinement-induced NE-cPLA2 signaling (Fig. 7).

**Fig. 7.**
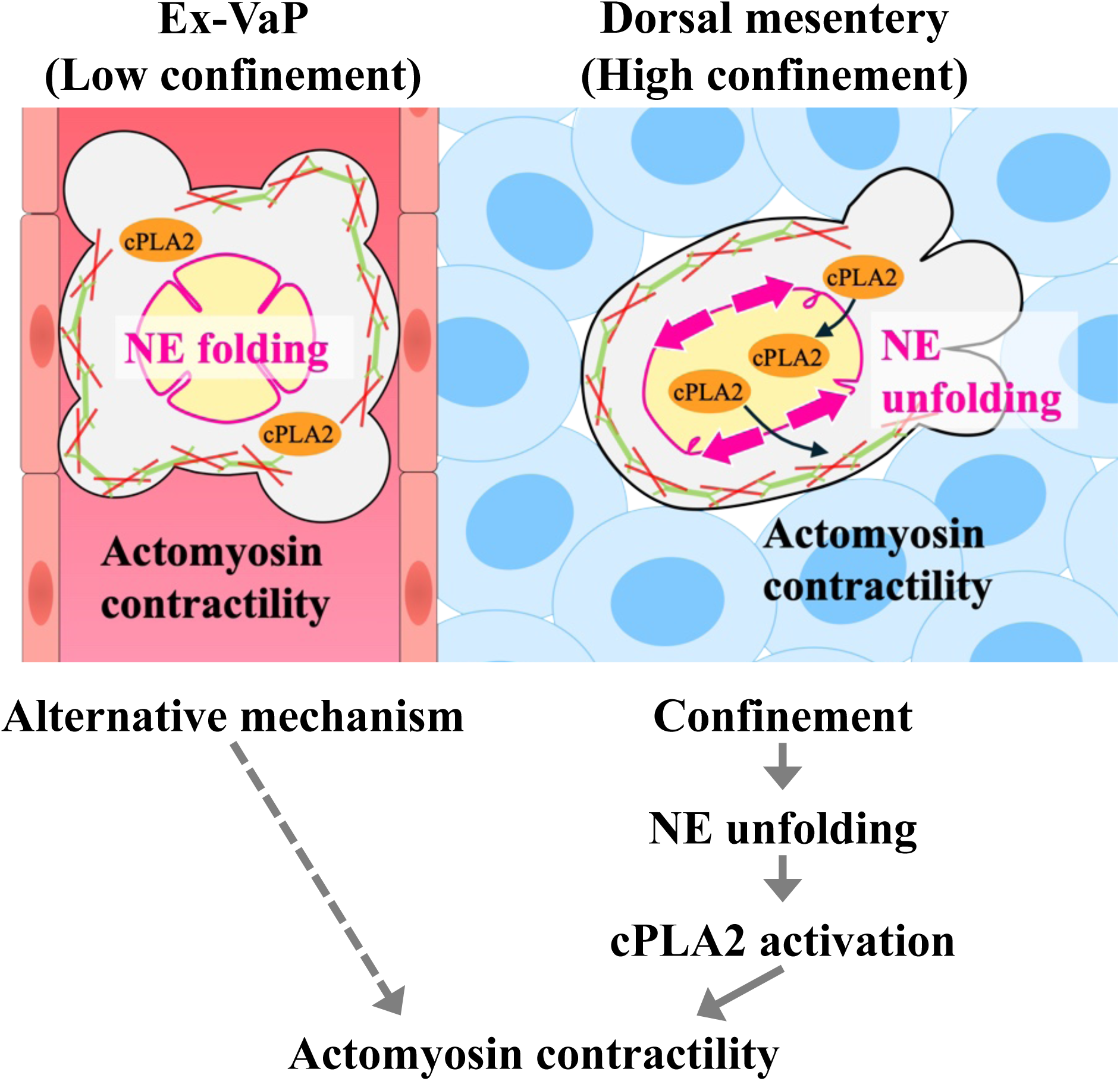
Model for environment-dependent regulation of bleb formation during PGC migration. PGCs employ a conserved bleb-based migratory mode in both the extravasation vascular plexus (Ex-VaP) and the dorsal mesentery (DM), where membrane blebs are generated by actomyosin contractility. In the DM, physical confinement imposed by surrounding mesenchymal cells (blue) induces nuclear envelope (NE) unfolding and nuclear accumulation of cPLA2, thereby promoting bleb formation. By contrast, PGCs crawling within the Ex-VaP generate membrane blebs independently of the confinement-induced NE-cPLA2 pathway, indicating that an alternative bleb-inducing mechanism operated in the vascular environment. These findings suggest that PGCs preserve a conserved bleb-based migratory program while switching the molecular mechanism of bleb formation according to the physical properties of their surrounding environment.

## DISCUSSION

One of the major findings of this study is the demonstration that the NE-cPLA2 pathway functions during physiological cell migration *in vivo*. Mechanical unfolding of the nuclear envelope has recently emerged as a mechanosensory mechanism linking physical confinement to membrane blebbing through activation of cPLA2 and cortical actomyosin contractility (Lomakin et al., 2020; Venturini et al., 2020). However, these studies relied largely on artificial confinement systems using cultured cells, leaving unresolved whether this pathway operates during physiological cell migration. Here, we show that PGCs migrating through the DM undergo NE unfolding, accompanied by nuclear accumulation of cPLA2 and cortical recruitment of myosin II, and that disruption of this pathway impairs both bleb formation and migration (Fig. 7). To our knowledge, this represents the first demonstration that the NE-cPLA2 pathway functions as a physiological mechanosensory module during embryonic cell migration.

Our previous study demonstrated that store-operated Ca²⁺ entry (SOCE) is essential for bleb formation during vascular crawling and transendothelial migration, as well as under *in vitro* confinement conditions resembling the DM (Morita et al., 2026). Together with the present findings, these results identify SOCE as a conserved signaling module required for bleb formation in both vascular and mesenchymal environments. In contrast, the present study demonstrates that the NE-cPLA2 pathway is selectively engaged only during migration through the DM, identifying it as an environment-specific bleb-inducing mechanism rather than a universal regulator of bleb formation. Whether the NE-cPLA2 pathway functions upstream of SOCE or operates in parallel with it remains unknown. Resolving this relationship will provide important insight into how a conserved bleb-forming machinery is differentially engaged in distinct tissue environments. Collectively, our findings support a hierarchical model of cell migration in which diverse environmental cues engage distinct signaling pathways that converge on a common SOCE-dependent bleb-forming machinery, thereby enabling a common bleb-based migratory program across heterogenous tissue environments.

An important unresolved question raised by this study concerns the physiological signals that initiate bleb formation during intravascular crawling. Unlike migration through the DM, where physical confinement activates the NE-cPLA2 pathway, vascular blebbing occurred independently of this signaling cascade, indicating that distinct regulatory mechanisms operate within the vascular environment. Several mechanical and biochemical cues represent plausible candidates for initiating this process. Our finding that vascular blebbing persists following experimental cessation of blood flow argues that continuous blood flow alone is unlikely to be sufficient for bleb induction (Extended Data Fig. 6a-d). Because cardiac arrest simultaneously alters multiple hemodynamic parameters, including shear stress, intravascular pressure, and pulsatile flow (Liu et al., 2011; Penfield and Montell, 2023), the contribution of each mechanical cue cannot be distinguished in the present study. Nevertheless, hydrostatic pressure generated as PGCs become lodged within narrow capillaries represents a plausible mechanical trigger (Chugh et al., 2022). Biochemical signaling may also contribute to vascular blebbing. We previously demonstrated that stem cell factor (SCF) is sufficient to induce persistent SOCE-dependent blebbing in cultured PGCs (Morita et al., 2026), raising the possibility that SCF produced within the Ex-VaP contributes to the initiation or maintenance of crawling blebs. Together, these findings identify both mechanical and biochemical signals as plausible upstream regulators of vascular blebbing while emphasizing that the physiological trigger for vascular blebbing remains unresolved.

Although this study focuses on avian primordial germ cells, the conceptual framework emerging from these findings may extend beyond the germline. Many migrating cell types, including immune cells, neural crest cells, and metastatic cancer cells, traverse multiple tissue environments with diverse mechanical properties (Yamada and Sixt, 2019). Our findings suggest that robust migration across heterogeneous tissues may not require fundamentally different migratory strategies but instead relies on flexible engagement of environment-specific signaling pathways that converge on a conserved migratory machinery responsible for generating productive protrusions. Such hierarchical regulation may represent a general principle by which migrating cells maintain a common migratory program while adapting to diverse physiological and pathological environments.

## MATERIALS and METHODS

### Animals, staging, and animal care

Fertilized chicken (*Gallus gallus domesticus*, White Leghorn) eggs were purchased from Yamagishi Poultry Farm (Mie, Japan). Fertilized eggs from wild-type Japanese quail (*Coturnix japonica*) and transgenic Japanese quail [TG(PGK1:H2B-chFP)] (Hess et al., 2015) were provided by Nagoya University through the National Bio-Resource Project of the MEXT, Japan. Eggs were incubated at 38.5℃, and embryos were staged according to the Hamburger and Hamilton (HH) staging system (Hamburger and Hamilton, 1951). All animal experiments were approved by the Institutional Animal Care and Use Committees at Kyushu University and were perfomed in accordance with institutional guidelines.

### PGC culture

Chicken PGCs were established from 1–2 μl of embryonic blood collected from HH15 White Leghorn embryos. Cells were maintained in FAcs medium at 38.0℃ under 5% CO_2_, as previously described (Chen et al., 2019; Whyte et al., 2015). PGCs were cultured in non-adherent 24-, 48-, or 96-well plates (Thermofisher Scientific), and half of the culture medium was replaced every other day.

FAcs medium consisted of 65.52% Ca^2+^-free DMEM (Nacalai tesque), 1× Nucleosides (Sigma-Aldrich), 2mM GlutaMAX^TM^ (Gibco), 1.2mM Sodium Pyruvate (Gibco), 1× non-essential amino acids (Gibco), 55 μM β-mercaptoethanol (Fujifilm Wako), 2 mg/ml Ovalbumin (Sigma-Aldrich), 1× penicillin-streptomycin-amphotericin B (Fujifilm Wako), 1× B-27 Supplement (Gibco), 0.2% chicken serum (Biowest), 0.1 mg/ml heparin sodium (Fujifilm Wako), 25ng/ml human Activin A (Apro Science), 4 ng/ml human FGF chimera (Fujifilm Wako), 100 µM CaCl_2_ (Nacalai tesque), 21.84% sterile distilled water (Fujifilm Wako).

### Plasmid constructions

Expression constructs were generated using the Tol2-based vectors pT2A-CAGGS or pT2A-BItight by conventional restriction enzyme cloning or In-Fusion HD cloning (TaKaRa). The coding sequences of Lifeact-mCherry, Tet3G, MRLC1-EGFP, mCherry-CAAX, LAP2b-AcGFP, LBR, ZsGreen1, mCherry, and cPLA2-EGFP were amplified by PCR or excised from existing plasmids and inserted into the appropriate vectors.

Tet-inducible constructs were generated using the bidirectional tetracycline-responsive vector pT2A-BItight-TRE. For co-expression experiments, LBR was combined with either LAP2b-AcGFP or ZsGreen1 in the same expression cassette. The cPLA2-EGFP construct was generated by In-Fusion assembly using PCR products amplified from chick embryonic cDNA and EGFP.

CRISPR/Cas9 vectors targeting the chicken PLA2g4a gene (NM_205423) were generated by designing sgRNAs using CHOPCHOP (https://chopchop.cbu.uib.no/) and cloning them into BbsI site of PX458 (Addgene plasmid #48138) (Ran et al., 2013).

RNAi expression vectors (pT2A-RNAiC-CAGGS-EGFP-IRES-PuroR and pT2A-RNAiC-CAGGS-mCherry-IRES-PuroR) were generated by amplifying the chicken miR-30 RNAi expression cassette from pRFPRNAi-C (Das et al., 2006) and cloning it into the SalI site of pT2A-CAGGS-EGFP-IRES-PuroR or pT2A-CAGGS-mCherry-IRES-PuroR. Target sequences were selected using siDirect (https://sidirect2.rnai.jp/) with a GC content of 30-60% and introduced into the miR-30 expression cassette by PCR.

Detailed cloning procedures, primer sequences, sgRNA sequences, and RNAi target sequences are provided in the Supplementary Methods.

### Genetic manipulation of PGCs

#### Plasmid transfection

Cultured PGCs were transfected by either electroporation or lipofection.

##### Electroporation

Electroporation was performed using the Neon Transfection System (Invitrogen). Briefly, 6×10^4^ PGCs were resuspended in Neon T buffer, mixed with plasmid DNA, and electroporated under optimized conditions (1,300 V, 10 ms, 3 pulses). Unless otherwise indicated, all expression constructs were co-transfected with the Tol2 transposase plasmid (pCAGGS-T2TP-pA) to achieve stable genomic integration. For Tet-inducible experiments, plasmids encoding the Tet-responsive construct, Tet3G transactivator, and Tol2 transposase were mixed at a ratio of 3:1:1. Cells were cultured overnight in antibiotic-free FAcs medium before replacement with standard FAcs medium.

##### Lipofection

For lipofection, 1 × 10^5^ PGCs were seeded into a non-adherent 96-well plate containing 150 µl of FAcs medium lacking heparin and antibiotics. A transfection mixture containing 1 µg plasmid DNA and 1 µl Lipofectamine 2000 (ThermoFisher Scientific) in 50 µl of Opti-MEM was incubated for 5 min at room temperature (RT) and then added to the cells. Unless otherwise indicated, all expression constructs were co-transfected with pCAGGS-T2TP-pA. The medium was replaced with standard FAcs medium 6 h after transfection.

For CRISPR-Cas9-mediated gene disruption, the PX458-sgRNA vectors targeting PLA2g4a described above were co-transfected into cultured PGCs. Complete disruption of PLA2g4a resulted in rapid cell death during culture, precluding subsequent migration analysis.

RNAi Knockdown efficiency was confirmed by RT-qPCR. Total RNA was was extracted using the NucleoSpin RNA XS kit (TaKaRa), and first-strand cDNA was synthesized using the PrimeScript II 1st Strand cDNA Synthesis Kit (TaKaRa). Quantitative PCR was performed using SYBR Green qPCR Master Mix (ThermoFisher Scientific) on a CFX Connect Real-Time PCR Detection System (Bio-Rad). PLA2g4a expression was normalized to β-actin. Primer sequences are provided in the Supplementary Methods.

#### Establishment of stable PGC lines

Stable transfectants carrying antibiotic resistance markers were selected in FAcs medium containing 0.5 µg/ml Puromycin (Gibco) for 2 days or 0.4 µg/ml G418 (FUJIFILM Wako) for 5 days, depending on the selection marker. Stable RNAi PGC lines were established by fluorescence-activated cell sorting (SH800, Sony) of EGFP- or mCherry-positive cells 5 days after transfection.

### *In ovo* transfection

*In ovo* transfection was performed as previously described (Saito et al., 2012). Lipofectamine 2000 (ThermoFisher Scientific) was diluted 1:1 in Opti-MEM (Gibco) and incubated for 5 min at RT. Plasmid DNA (pT2A-CAGGS-Lifeact-mCherry) was diluted in Opti-MEM to a final concentration of 250 μg/ml. Equal volumes of the Lipofectamine and DNA solutions were then mixed and incubated for 20 min at RT to allow complex formation. A total of 1 µl of the transfection mixture was injected into the bloodstream of HH15 chicken embryos.

#### PGC transplantation and Tet induction

Cultured PGCs were washed with Opti-MEM and resuspended at 5×10^3^ cells/µl in Opti-MEM. A total of 5×10^3^ PGCs (1 µl) were injected into the heart of HH15 embryos using a glass capillary (GD-1, NARISHIGE) pulled with a Puller PC-100 micropipette puller (NARISHIGE).

For induction of Tet-inducible constructs *in vitro*, PGCs were cultured in FAcs medium containing 1 µg/ml doxycycline (Dox) (Clontech) for either 12 hours or 3 days before transplantation. For *in ovo* induction, 500 µl of 50 µg/ml Dox (prepared by diluting a 1 mg/ml stock solution 1:20 in Hanks’ balanced salt solution) was applied onto the embryo immediately after transplantation. Embryos were subsequently incubated at 38.5℃ until the desired developmental stage.

#### Pharmacological treatments

For pharmacological inhibition, cultured PGCs were seeded into 24-well plates containing 500 µl FAcs medium and treated with either 100 µM blebbistatin (Fujifilm Wako) for 1 h or 25 µM AACOCF3 (Cayman Chemical) for 2 h. Following treatment, cells were washed once with Opti-MEM (Gibco) and returned to fresh FAcs medium for subsequent experiments.

#### Live imaging preparations

##### Whole-mount *ex vivo* live imaging

Whole-mount ex vivo live imaging was performed as previously described (Saito et al., 2022), HH15 chicken embryos transplanted with cultured PGCs were prepared as described above. For visualization of the embryonic vasculature during live imaging, Alexa Fluor 488- or Alexa Fluor 594-conjugated acetylated low-density lipoprotein (ThermoFisher Scientific, L23380 and L35353) was diluted 1:2 in Opti-MEM and co-injected with PGC suspension. Glass-bottom dishes (50 mm; Matsunami) were coated with a thin layer of albumen collected from unincubated eggs and incubated at 38.5℃. Immediately before embryo culture, excess albumen was removed. Embryos were carefully transferred onto the coated dishes with the ventral side facing the glass surface and incubated for 10 min at 38.5℃ to allow attachment. Additional thin albumen was then added to cover the embryos, and samples were maintained at 38.5℃ throughout live imaging.

##### *Ex vivo* live imaging of dissected dorsal mesentery

Glass bottom dishes (35 mm; Eppendorf) were coated with 10 µg/ml poly-L-lysine hydrochloride (Peptide Institute) for 1 h at RT, washed three times with distilled water, and air-dried. The culture medium consisted of DMEM (08458-45, Nacalai tesque), supplemented with 10% fetal bovine serum, 2 mM GlutaMax (Gibco), and 1× penicillin-streptomycin-amphotericin B (FUJIFILM Wako). To immobilize the tissue during imaging, 2% low-melting-point agarose (Promega) dissolved in PBS containing 0.3% glucose was mixed immediately before use with an equal volume of pre-warmed culture medium. 12 hours after PGC transplantation, embryos were removed from the eggs and washed in PBS and cleared of extra-embryonic membranes. The dorsal mesentery was dissected, placed ventral side down on the poly-L-lysine-coated surface with a minimal volume of culture medium, overlaid with 150 µl of the agarose-medium mixture, and allowed to solidify for 1 min at 4℃. Culture medium was then added, and samples were maintained at 38℃ under 5% CO_2_ until imaging.

##### Under agarose gel assay

The under -agarose confinement assay was performed as previously described (Heit and Kubes, 2003; Kirkness et al., 2018; Morita et al., 2026), with minor modifications. To prevent cell adhesion, 35-mm glass-bottom dishes (Eppendorf) were coated with 10 mg/ml Pluronic F127 (Sigma-Aldrich) for 30 min at RT, and then air-dried. Agarose gels were prepared by mixing 2x Hank’s balanced salt solution (HBSS) (Gibco) and FAcs medium at a ratio of 1:2. UltraPure agarose (Invitrogen) was dissolved in distilled water at 48 mg/ml, and the agarose solution was mixed with the HBSS/FAcs solution at a ratio of 1:5. A total of 3ml of the mixture was added to each dish and allowed to solidify at RT. Gels were equilibrated with 500 µl of FAcs medium at 38℃ under 5% CO_2_ for 30 min. For confinement experiments, 1× 10^5^ cultured PGCs were suspended in 10 µl of FAcs medium containing polystyrene microbeads (Sigma-Aldrich) with diameters of 10 µm or 7µm. After removal of the agarose gel, the cell suspension was applied to the glass surface, the gel was replaced to confine the cells, and 50 μl of FAcs medium was added. Samples were incubated at 38℃ under 5% CO_2_ before live imaging.

#### FLIM measurements of plasma membrane tension

Plasma membrane tension was quantified using the fluorescent membrane tension probe Flipper-TR (Spirochrome, SC020) as previously described (Colom et al., 2018). For analysis of cultured PGCs, cells were incubated in serum-free FAs medium containing 1 µM Flipper-TR for 1 h. For analysis of PGCs migrating in the DM, dissected quail dorsal mesenteries were incubated in 2 µM Flipper-TR for 2 h. Transgenic quail Tg (PGK1:H2B-chFP) (Huss et al., 2015) were used to identify PGCs by nuclear fluorescence. For imaging under unconfined conditions, stained PGCs were suspended in 10% Matrigel Basement Membrane Matrix Growth Factor Reduced, Phenol Red Free (Corning) to prevent cell displacement during image acquisition. For confinement experiments, PGCs were prepared using the under-agarose confinement assay described above. FLIM images were acquired using a STELLARIS 8 FALCON microscope (Leica) equipped with an HC PL APO 63x/1.20 W CORR CS2 objective and Las X software. Flipper-TR was excited at 488 nm using a pulsed white-light laser operating at 80 MHz, and fluorescence emission was collected between 550 and 650 nm using a HyD X detector with 16-bit digitization and a pinhole set to 1 Airy unit. Samples were maintained at 38℃ under 5% CO_2_ using an incubation chamber (TOKAI HIT STX) throughout image acquisition. FLIM image processing and fluorescence lifetime analysis were performed as previously described (Wang et al., 2023), with minor modifications. Background fluorescence was excluded by applying an intensity threshold of 200 photons per pixel, and plasma membrane regions were manually selected as regions of interest for fluorescence lifetime measurements,

#### Experimental cessation of blood flow in Ex-VaP

To examine whether blood flow is required for bleb formation during vascular crawling, blood flow was experimentally arrested by inhibiting cardiac contraction. 1 h after PGC transplantation, embryos were prepared for whole-mount *ex vivo* live imaging as described above. Cardiac contraction was inhibited by applying 100 mM 2,3-butanedione-2-monoxime (BDM; Cayman Chemical) (Banjo et al., 2013) dissolved in PBS directly onto the embryonic heart. To confirm that transient BDM treatment did not impair embryonic development, somite numbers were counted immediately before and 3 h after treatment. No significant difference in somite number was observed between BDM-treated and control embryos.

To verify cessation of blood flow, 1 µl of FluoSpheres polystyrene microspheres (Invitrogen) was injected into the circulation, and microsphere movement within the Ex-VaP was recorded using an MVX10 stereomicroscope (Olympus) equipped with an ORCA-R2 C10600-10B camera (Hamamatsu Photonics). PGC behavior within the Ex-VaP was imaged immediately after BDM treatment.

#### Fixation, whole-mount, and section immunostaining

Chicken and quail embryos were fixed in 4% paraformaldehyde (PFA) in PBS for 16-24 h at 4℃. Samples were cryoprotected in 30% sucrose/PBS for 3-6 h, incubated overnight at 4℃ in a 2:1 mixture of OCT compound (Tissue-Tek) and 30% sucrose/PBS, embedded in OCT compound, and sectioned at 14-µm using a CryoStar NX70 cryostat (ThermoFisher Scientific). Cryosections were washed three times for 5 min in TNT buffer (0.1 M Tris-HCl pH 7.5, 0.15 M NaCl, 0.05% Tween 20) and blocked for 1 h at RT in 1% Blocking Reagent (Roche) prepared in TNT buffer.

Primary antibodies used in this study included goat anti-EGFP (BIO-RAD, 1:1,000), rabbit anti-RFP (Funakoshi, 1:1,000), and mouse QH1 (DSHB, 1:40) and a rabbit anti-DDX4 antibody (Eurofins Genomics; 1:1,000) raised against the N-terminal peptide (MEEDWDTELEQE) of chicken DDX4 (Saito et al., 2022).

Secondary antibodies included Alexa Fluor 488-conjugated donkey anti-goat IgG (Invitrogen, A11055), Alexa Fluor 555-conjugated donkey anti-rabbit IgG (Invitrogen, A31572), and Alexa Fluor 647-conjugated donkey anti-mouse IgG (Invitrogen, A31571), all used at a dilution of 1:500. F-actin was visualized using Alexa Fluor 568- or Alexa Fluor 647-conjugated Phalloidin (Invitrogen, 1:100), as indicated. Finally, sections were mounted in DAPI Fluoromount-G (SouthernBiotech).

#### Microscopy

Unless otherwise specified, confocal fluorescence images were acquired using a Dragonfly high-speed spinning-disk confocal microscope (Oxford Instruments). Three-dimensional image reconstruction and rendering were performed using Imaris software (Oxford Instruments). Imaging systems used FLIM and blood flow measurements are described in the corresponding sections.

#### Quantitative image analysis

##### Quantification of blebbing PGCs

To quantify bleb formation, cells exhibiting at least one cycle of bleb expansion and retraction during time-lapse imaging were scored as blebbing PGCs. Images were acquired every 30 s for 30 min during whole-mount *ex vivo* imaging and every 20 s for 2 min during the under-agarose confinement assay. The percentage of blebbing PGCs was calculated relative to the total number of analyzed cells.

##### Nuclear envelope invagination ratio

The nuclear envelope (NE) invagination ratio was quantified as previously described (Venturini et al., 2020). Confocal z-section images of LAP2b-AcGFP-expressing PGCs were analyzed using Fiji (ImageJ). The contour of the inner nuclear membrane was manually traced, and both the enclosed area (A) and the corresponding convex hull area (C) were measured. The NE invagination ratio was calculated as 1 - (A/C), with higher values indicating a greater degree of NE invagination (Venturini et al., 2020).

##### Nuclear envelope wrinkle index

The nuclear envelope (NE) wrinkling index was quantified as previously described (Cosgrove et al., 2021) with minor modifications. Maximum-intensity projections of confocal z-stacks of LAP2b-AcGFP-expressing PGCs were generated using Fiji (ImageJ). The mean fluorescence intensity (μ) and standard deviation (σ) of the nuclear signal were measured, and the threshold for edge detection using FeatureJ Edges plugin (Sobel operator) was calculated as (μ − 1.5σ)/Brf, where Brf denotes the brightness regularization factor. To optimize edge detection while minimizing background signals, a range of Brf and smoothing scales was evaluated. A Brf value of 8 and a smoothing scale of 1.5 were used for all analyses. The NE wrinkle index was calculated as the mean intensity of the detected edge signal divided by the maximum pixel intensity (255) and multiplied by 100.

##### Cortical myosin II enrichment

Cortical myosin II enrichment was quantified as previously described (Ruprecht et al., 2015). Confocal z-stack images of MRLC1-EGFP-expressing PGCs were analyzed using Fiji (ImageJ). Fluorescence intensity line profiles (20-pixel width) were drawn across the cell cortex. The average cytoplasmic fluorescence intensity (F_cyto_) and the maximum cortical fluorescence intensity (F_max_) were determined from each line profile, where F_max_ was defined as the mean of the three highest intensity values. Cortical myosin II enrichment was calculated as (F_max_-F_cyto_)/F_cyto_.

### Statistical analysis

Data are presented as mean ± SD, unless otherwise indicated. Statistical analyses were performed using GraphPad Prism 10 (GraphPad Software). For comparisons between two groups, two-sided unpaired Student’s t-tests were used. For comparisons among three or more groups, one-way analysis of variance (ANOVA) followed by Tukey’s post hoc test was performed. Statistical tests, sample sizes (*n*), and the numbers of independent experiments are indicated in the corresponding figure legends. Differences were considered statistically significant at P < 0.05.

## ACKNOWLEDGEMENTS

We thank the Center for Advanced Instrumental and Educational Support of the Faculty of Agriculture, Kyushu University, for access to shared research facilities, including fluorescence-activated cell sorting (SH800), cryostat sectioning, and confocal microscopy (STELLARIS). We thank the National BioResource Project (NBRP), Japan, and the Graduate School of Bioagricultural Sciences, Nagoya University, for providing quail eggs and transgenic quail lines. We thank the Imaging Facility of the National Institute for Basic Biology (NIBB) for technical assistance with fluorescence lifetime imaging microscopy (FLIM). This work was also supported by the NIBB Cooperative Research Program (#25NIBB535 to D. S.) and the Advanced Bioimaging Support (ABiS) platform (JSPS KAKENHI Grant Number JP22H04926). This work was supported by JSPS KAKENHI (Grant numbers 23K23897 and 26K02019 to D. S.), a JSPS Research Fellowship for Young Scientists (DC2; Grant Number JP26KJ1802 to M. Morimoto), JST SPRING (Grant Number JPMJSP2136 to M. Morimoto), the Shinnihon Foundation of Advanced Medical Treatment Research, and the Terumo Life Science Foundation (to D. S.).

## AUTHOR CONTRIBUTIONS

M. Morimoto and D. S. designed the study. M. Morimoto performed most experiments and analyzed data. M. Morita performed the live imaging experiments. Y. K. performed FLIM image acquisition. D. S., J. I., and Y. H. supervised the study. M. Morimoto and D. S. wrote the manuscript.

## DECLARATION OF INTERESTS

The authors declare no competing interests.

**Extended Data Figure 1. Endogenous PGCs exhibit membrane blebs in Ex-VaP and the dorsal mesentery.**

**a,** Representative image of an endogenous PGC crawling within the Ex-VaP stained with *Lycopersicon Esculentum* (Tomato) lectin (LEL). Arrows indicate bleb-like protrusions. **b,** Schematic of *in ovo* transfection of Lifeact-mCherry plasmid DNA into HH15 chicken embryos and analysis of transverse sections at HH19 (left). Representative image of an endogenous PGC at HH19 stained for DDX4 (green) and mCherry (Red). Arrowheads indicate membrane blebs lacking cortical F-action. Scale bars, 10 µm.

**Extended Data Figure 2. Under-agarose confinement assay with defined confinement heights.**

**a,** Schematic of the under-agarose confinement assay. PGCs were placed between a glass coverslip and an agarose gel pad containing microbeads, which defined the confinement height according to bead diameter (10 or 7 µm). **b**, Representative confocal lateral views of PGCs expressing Lifeact-mCherry and LAP2b-AcGFP under unconfined conditions or under 10-µm or 7-µm confinement. The resulting cell height was approximately 15-20 µm under unconfined conditions, 8-10 µm under 10-µm confinement, and 5-7 µm under 7-µm confinement. Scale bars, 5 µm.

**Extended Data Figure 3. LBR overexpression does not affect PGC proliferation or arrest within the Ex-VaP.**

**a,** Cell numbers of control (mCherry or ZsGreen1) and LBR-OE (ZsGreen+LBR) PGCs after 5 days of culture initiated from 1,000 cells (n = 5 independent cultures). **b,c,** Representative whole-mount images (b) and quantification (c) of transplanted control and LBR-OE PGCs arrested within the Ex-VaP at HH17. The ratio of ZsGreen1-positive (control or LBR-OE) to co-transplanted mCherry-positive PGCs was quantified (Control: n = 4 embryos; LBR-OE: n = 5 embryos). Scale bars, 100 µm. n.s., not significant (two-sided unpaired Student’s t-test).

**Extended Data Figure 4. Combined cPLA2 knockdown and AACOCF3 treatment suppresses bleb formation.**

**a**, Relative cPLA2g4a mRNA expression in PGCs expressing si-Ctrl (EGFP), si-Ctrl (mCherry), or si-cPLA2. Expression levels were normalized to si-Ctrl (EGFP) (n = 5 independent experiments; 1×10^5^ cells per experiment). **b,c,** Representative images (b) and quantification (c) of bleb formation in si-Ctrl and si-cPLA2 PGCs under 10-µm confinement. Data are presented as mean ± SD from four independent experiments (n = 4), with approximately 100 cells analyzed per experiment. **d,e,** Representative images (d) and quantification (e) of bleb formation in si-Ctrl+EtOH, si-Ctrl+AACOCF3 (AA), and si-cPLA2+AA PGCs under 10-µm confinement. Data are presented as mean ± SD from three independent experiments (n = 3), with approximately 100 cells analyzed per experiment. Scale bars, 10 µm. n.s., not significant. *P<0.05, **P<0.01 (two-sided unpaired Student’s t-test).

**Extended Data Figure 5. Experimental design and validation of the cPLA2 inhibition assay for *in vivo* PGC migration.**

**a,** After 2 h of treatment with vehicle (ethanol) or AACOCF3 (AA), equal numbers of EGFP-labeled si-Ctrl+EtOH or si-cPLA2+AA PGCs were co-transplanted with mCherry-labeled control PGCs into the circulation of HH15 chick embryos. At HH21 (1 day after transplantation), EGFP-positive and mCherry-positive PGCs within the developing gonads were quantified. **b,** Cell number of si-Ctrl+EtOH and si-cPLA2+AA PGCs 24 h after drug treatment. Cells were reseeded at 50,000 cells immediately after treatment (n = 3 independent experiments). n.s., not significant. (two-sided unpaired Student’s t-test).

**Extended Data Figure 6. PGCs continue bleb-based crawling after experimental cessation of blood flow in the Ex-VaP.**

**a,** Schematic of the experimental design for experimental cessation of blood flow following PGC transplantation. **b**, Blood flow velocity within the Ex-VaP in PBS-treated (control) and BDM-treated embryos (n = 3 embryos per condition). **c**, Representative *ex vivo* live images of transplanted PGCs crawling within the Ex-VaP in PBS-treated (control) and BDM-treated embryos. Arrowheads indicate membrane blebs. **d**, Quantification of the proportion of blebbing PGCs crawling within the Ex-VaP in PBS-treated (control) and BDM-treated embryos (n = 6 embryos per condition). Scale bars, 10 µm. n.s., not significant. *P<0.05 (two-sided unpaired Student’s t-test).

## VIDEO CAPTIONS

**Supplementary Videos1∼**

**作成中**

