## Supplemental Figures for "Environment-specific mechanosensing preserves a common bleb-based migratory program across diverse embryonic environments"

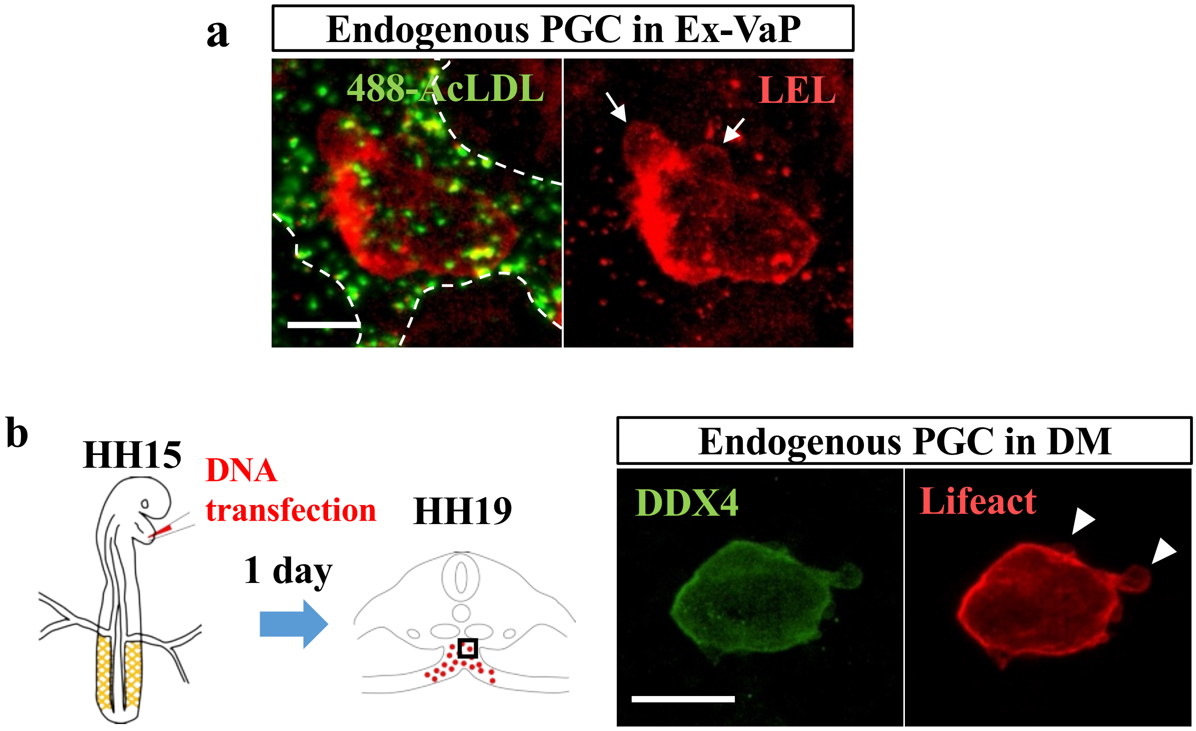


**Extended Data Figure 1 Morimoto *et al.***

**Extended Data Figure 1.**
**Endogenous PGCs exhibit membrane blebs in Ex-VaP and the dorsal mesentery.**
**a,** Representative image of an endogenous PGC crawling within the Ex-VaP stained with *Lycopersicon Esculentum* (Tomato) lectin (LEL). Arrows indicate bleb-like protrusions. **b,** Schematic of *in ovo* transfection of Lifeact-mCherry plasmid DNA into HH15 chicken embryos and analysis of transverse sections at HH19 (left). Representative image of an endogenous PGC at HH19 stained for DDX4 (green) and mCherry (Red). Arrowheads indicate membrane blebs lacking cortical F-action. Scale bars, 10 μm.

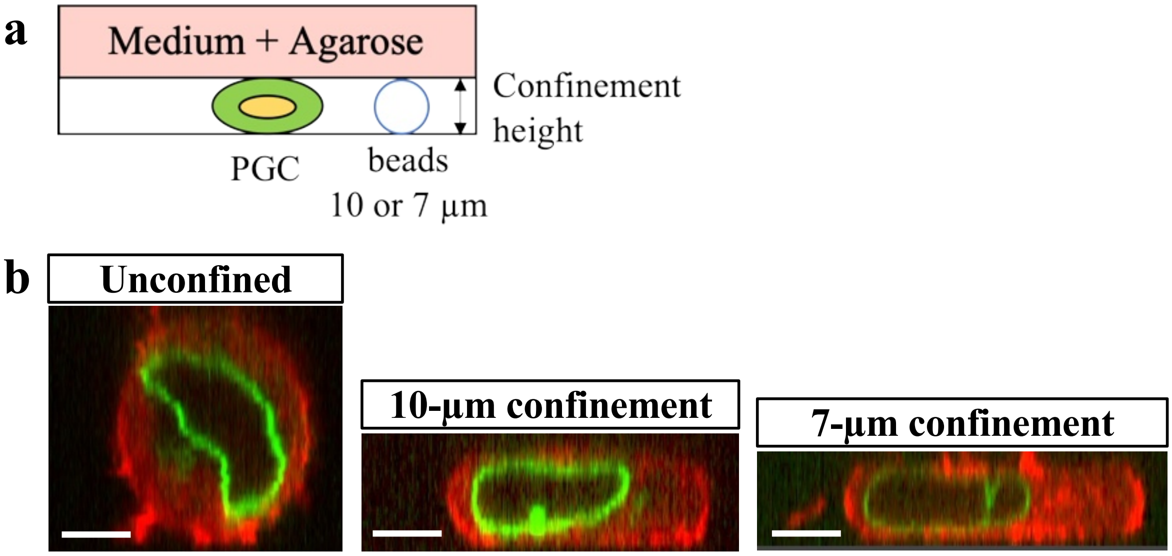


**Extended Data Figure 2 Morimoto *et al.***

**Extended Data Figure 2.**
**Under-agarose confinement assay with defined confinement heights.**
**a,** Schematic of the under-agarose confinement assay. PGCs were placed between a glass coverslip and an agarose gel pad containing microbeads, which defined the confinement height according to bead diameter (10 or 7 µm). **b**, Representative confocal lateral views of PGCs expressing Lifeact-mCherry and LAP2b-AcGFP under unconfined conditions or under 10-µm or 7-µm confinement. The resulting cell height was approximately 15-20 µm under unconfined conditions, 8-10 µm under 10-µm confinement, and 5-7 µm under 7-µm confinement. Scale bars, 5 µm.

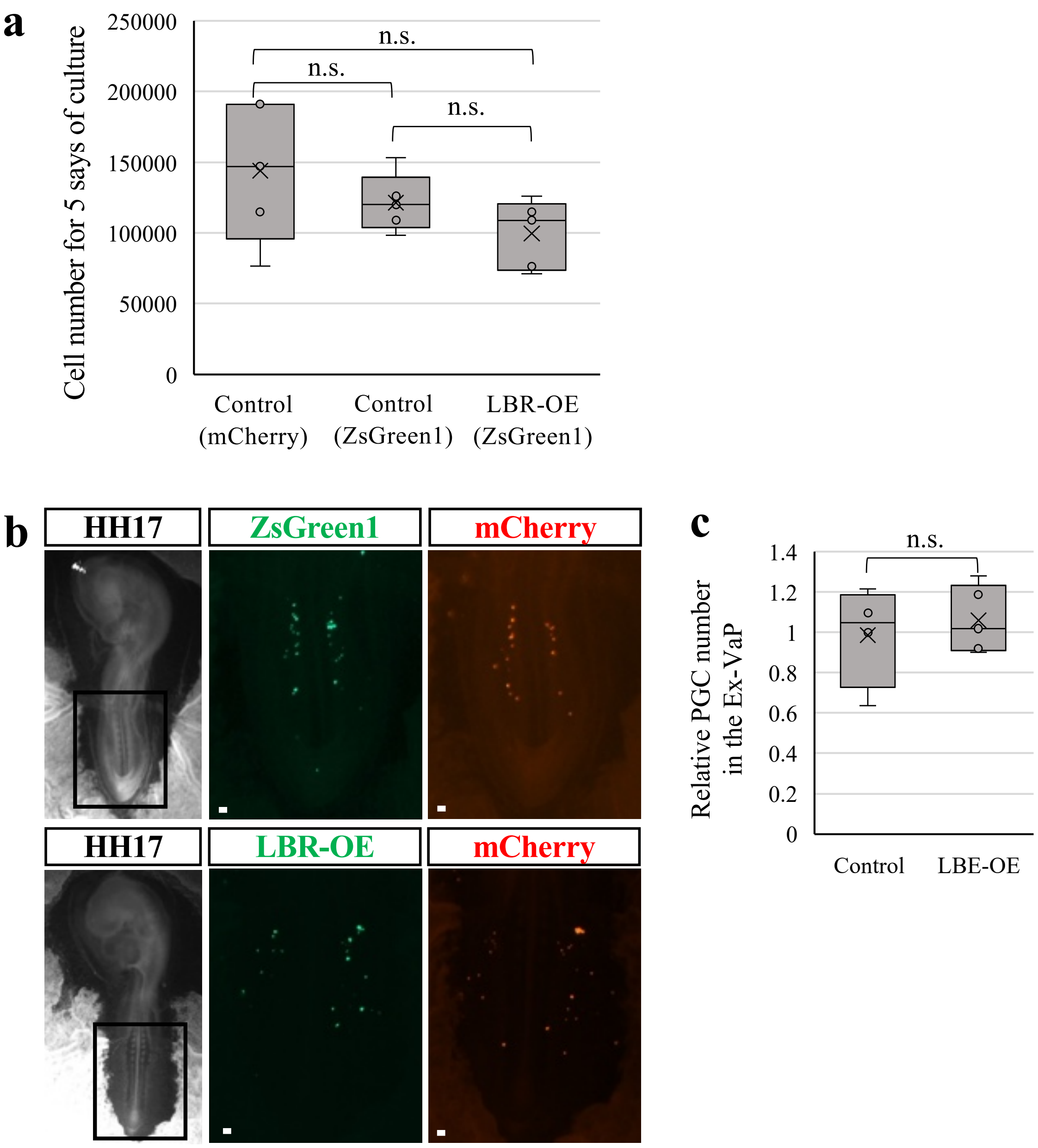


**Extended Data Figure 3 Morimoto *et al.***

**Extended Data Figure 3.**

**LBR overexpression does not affect PGC proliferation or arrest within the Ex-VaP.**

**a,** Cell numbers of control (mCherry or ZsGreen1) and LBR-OE (ZsGreen+LBR) PGCs after 5 days of culture initiated from 1,000 cells (n = 5 independent cultures). **b,c,** Representative whole-mount images (b) and quantification (c) of transplanted control and LBR-OE PGCs arrested within the Ex-VaP at HH17. The ratio of ZsGreen1-positive (control or LBR-OE) to co-transplanted mCherry-positive PGCs was quantified (Control: n = 4 embryos; LBR-OE: n = 5 embryos). Scale bars, 100 µm. P-values calculated by one-way ANOVA (a) and two-sided unpaired Student's t-test (c). n.s., not significant.

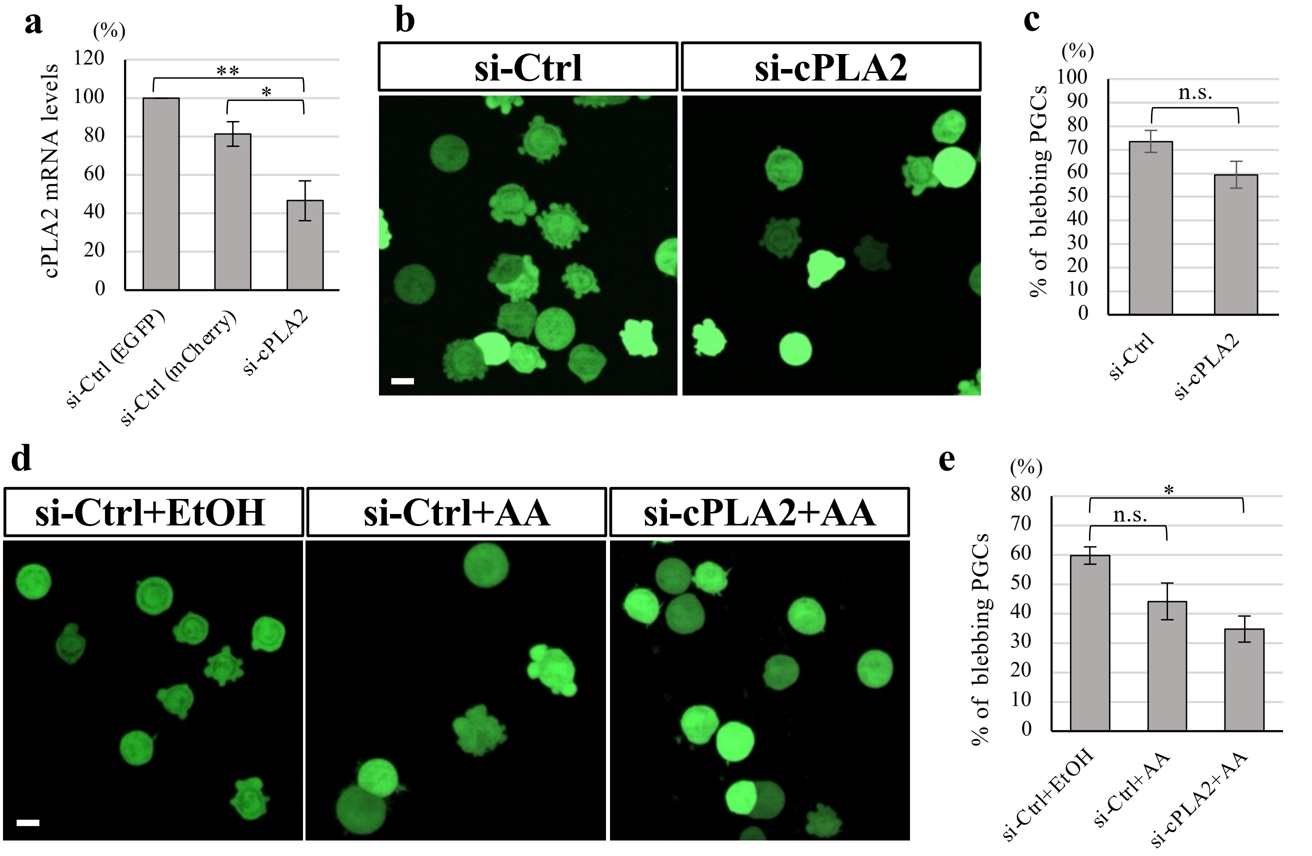


**Extended Data Figure 4 Morimoto *et al.***

**Extended Data Figure 4.**

**Combined cPLA2 knockdown and AACOCF3 treatment suppresses bleb formation.**

**a**, Relative cPLA2g4a mRNA expression in PGCs expressing si-Ctrl (EGFP), si-Ctrl (mCherry), or si-cPLA2. Expression levels were normalized to si-Ctrl (EGFP) (n = 5 independent experiments; $1 \times 10^5$ cells per experiment). **b,c,** Representative images (b) and quantification (c) of bleb formation in si-Ctrl and si-cPLA2 PGCs under 10-µm confinement. Data are presented as mean ± SD from four independent experiments (n = 4), with approximately 100 cells analyzed per experiment. **d,e,** Representative images (d) and quantification (e) of bleb formation in si-Ctrl+EtOH, si-Ctrl+AACOCF3 (AA), and si-cPLA2+AA PGCs under 10-µm confinement. Data are presented as mean ± SD from three independent experiments (n = 3), with approximately 100 cells analyzed per experiment. Scale bars, 10 µm. P-values calculated by one-way ANOVA (a,e) and two-sided unpaired Student's t-test (c). n.s., not significant. $*P<0.05$, $**P<0.01$.

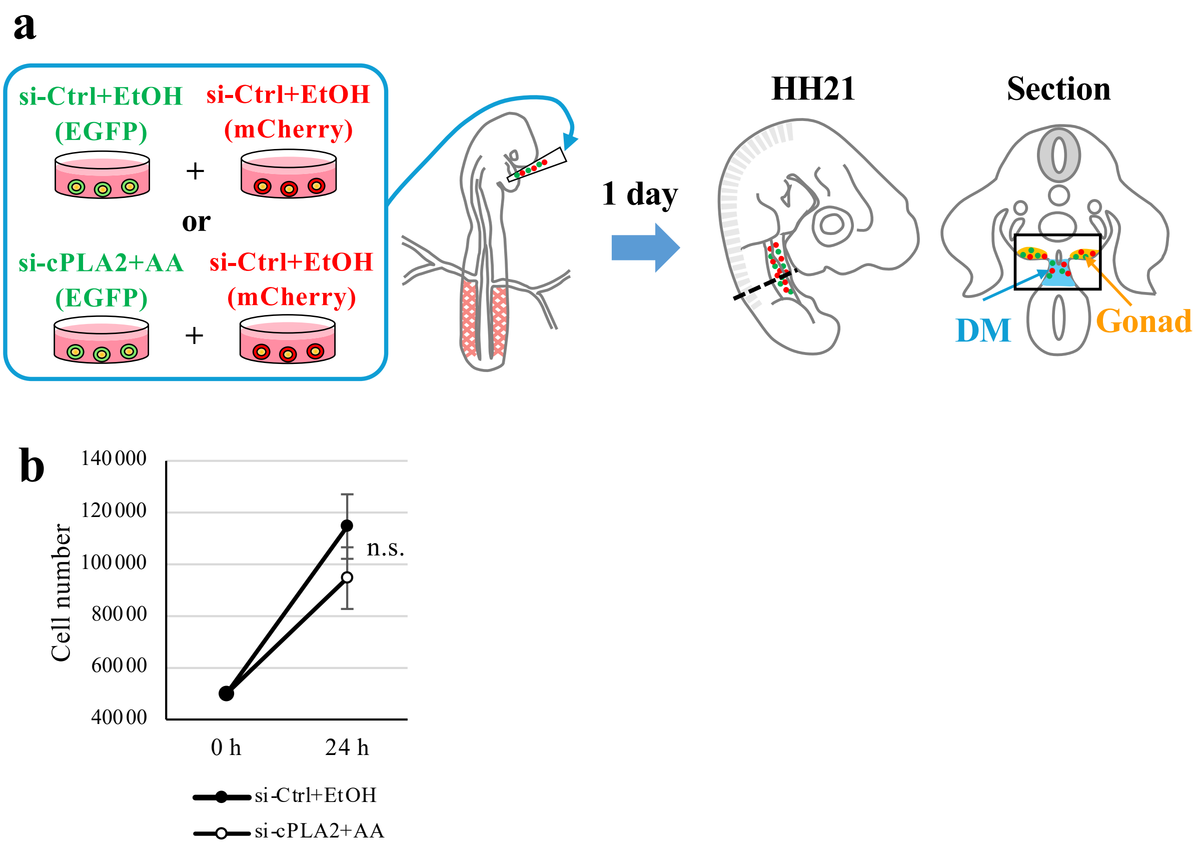


Extended Data Figure 5 Morimoto *et al.*

**Extended Data Figure 5.**
**Experimental design and validation of the cPLA2 inhibition assay for *in vivo* PGC migration.**
**a,** After 2 h of treatment with vehicle (ethanol) or AACOCF3 (AA), equal numbers of EGFP-labeled si-Ctrl+EtOH or si-cPLA2+AA PGCs were co-transplanted with mCherry-labeled control PGCs into the circulation of HH15 chick embryos. At HH21 (1 day after transplantation), EGFP-positive and mCherry-positive PGCs within the developing gonads were quantified. **b,** Cell number of si-Ctrl+EtOH and si-cPLA2+AA PGCs 24 h after drug treatment. Cells were reseeded at 50,000 cells immediately after treatment (n = 3 independent experiments). n.s., not significant (two-sided unpaired Student’s t-test).

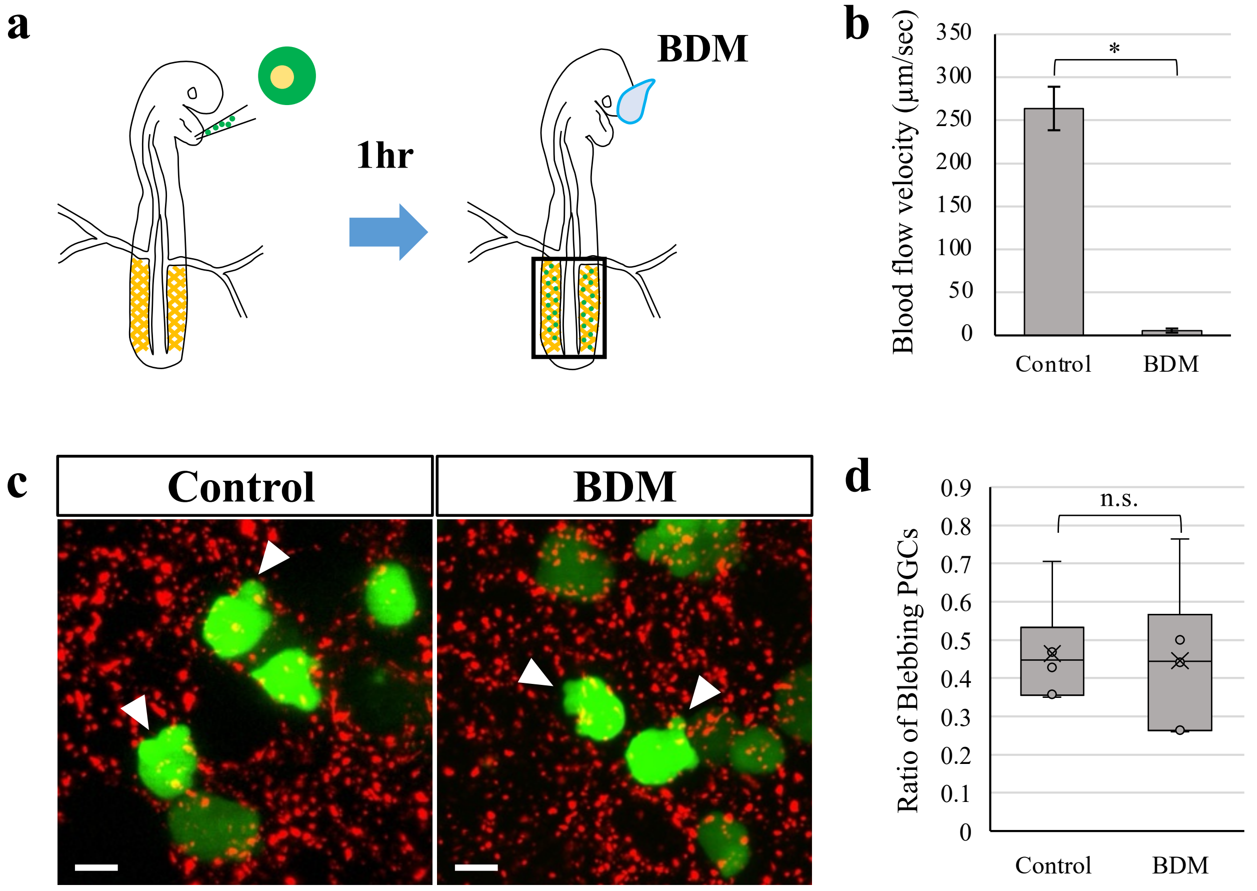


**Extended Data Figure 6 Morimoto *et al.***

**Extended Data Figure 6.**

**PGCs continue bleb-based crawling after experimental cessation of blood flow in the Ex-VaP.**

**a,** Schematic of the experimental design for experimental cessation of blood flow following PGC transplantation. **b**, Blood flow velocity within the Ex-VaP in PBS-treated (control) and BDM-treated embryos (n = 3 embryos per condition). **c**, Representative *ex vivo* live images of transplanted PGCs crawling within the Ex-VaP in PBS-treated (control) and BDM-treated embryos. Arrowheads indicate membrane blebs. **d**, Quantification of the proportion of blebbing PGCs crawling within the Ex-VaP in PBS-treated (control) and BDM-treated embryos (n = 6 embryos per condition). Scale bars, 10 µm. P-values calculated by two-sided unpaired Student's t-test (b,d). n.s., not significant. *$P<0.05$.
