## Supplemental Methods for "Environment-specific mechanosensing preserves a common bleb-based migratory program across diverse embryonic environments"

**Supplementary method**

Plasmid constructions

| Construct | Primer name | Sequence | Restriction site |
|---|---|---|---|
| pT2A-CAGGS-Tet3G-IRES2-PuroR | Tet3G-NotI-F | gcagcggccgcATGTCTAGACTGGACAAGAGCAAAGTC | NotI-XhoI |
| | Tet3G-XhoI-R | ggcctcgagTTACCCGGGGAGCATGTCAAGGTCAAAAT | |
| pT2A-BItight-Lifeact-mCherry-TRE | None | None | XhoI-BglII |
| pT2A-CAGGS-MRLC1-EGFP-IRES-P | MRLC1-NotI-F | tgcagcggccgcATGTCCAGCAAGCGGGCCAA | NotI-XhoI |
| | EGFP-XhoI-R | gggcctcgagTTACTTGTACAGCTCGTCCATG | |
| pT2A-CAGGS-mCherry-CAAX-IRES-PuroR | None | None | NotI-MluI |
| pT2A-BItight-LAP2b-AcGFP-TRE | LAP2b- MluI-F | tcctctagtcagctgacgcgtATGCCGGAGTTCCTAGAGGACCCTTCGG | MluI-EcoRV |
| | AcGFP-EcoRV-R | ttatgatcctctggagatatcTCACTTGTACAGCTCATCCATGCCGTGG | |
| pT2A-BItight-LBR-TRE-LAP2b-AcGFP | LBR-EcoRI-F | gtcagatcgcctggagaattcATGCCAAACCGGAAGTATGCCGATG | EcoRI-BglII |
| | LBR-BglII-R | gactgcagcctcaggagatctCTAGTAGATGTAAGGAAATATGCGGTAT | |
| pT2A-BItight-ZsGreen1-TRE | ZsGreen-EcoRI-F | ggagaattcATGGCCCAGTCCAAGCACGGCCTGACCAA | EcoRI-PstI |
| | ZsGreen-PstI-R | agactgcagTTAGGGCAAGGCGGAGCCGGAGGCGATGG | |
| pT2A-BItight-ZsGreen1-TRE-LBR | LBR-NotI-F | gacgcggccgcATGCCAAACCGGAAGTATGCCGATGGCGAGGTG | NotI-MulI |
| | LBR-MluI-R | agcacgcgtCTAGTAGATGTAAGGAAATATGCGGTATGGTACACGC | |
| pT2A-BItight-mCherry-TRE | mCherry-EcoRI-F | ggagaattcATGGTGAGCAAGGGCGAGGAGGATAACATGG | EcoRI-BglII |
| | mCherry-BglII-R | caggagatctTTACTTGTACAGCTCGTCCATGCCGCCGGT | |
| pT2A-BItight-TRE-cPLA2-EGFP | cPLA2-MluI-F | cctctagtcagctgacgcgtATGTCCTTCATAGATCCTTATCAACAC | MluI-NotI |
| | cPLA2-linker-R | CAAGAACAATCTTAACAACCATACGggagctggaggaattcgtcctgg | |
| | EGFP-linker-F | ctggaggaattcgtcctggatctATGGTGAGCAAGGGCGAGGAGCTGTTC | |
| | EGFP-NotI-R | cttatcgatgcggccgcTTACTTGTACAGCTCGTCCATGCCGAG | |

PLA2g4a knockout target sequences

| Name | Sequence |
|---|---|
| cPLA2-KO1 | ATACTCACGCATGTCGCCAATGG |
| cPLA2-KO2 | ATTCATTCTGGACCCCAACCAGG |

Target sequence for RNAi

| Name | Sequence |
|---|---|
| Scramble control (SC) | GTCATGAAAGCGCTTTATGAAT |
| cPLA2 | CCATACGAATCAGTAATGAATG |

Primers for RNAi hairpin

| Name | Sequence |
|---|---|
| First hairpin primer 5 | ggcggggctagctggagaagatgccttccggagaggtgctgctgagcg |
| First hairpin primer 3 | gggtggacgcgtaagaggggaagaaagcttctaaccccgctattcaccaccactaggca |
| SC forward | gagaggtgctgctgagcgaCATACGAATCAGTAATGAATGtagtgaagccacagatgta |
| SC reverse | attcaccaccactaggcaCCATACGAATCAGTAATGAATGtacatctgtggcttcact |
| cPLA2 forward | gagaggtgctgctgagcgtTCATGAAAGCGCTTTATGAATtagtgaagccacagatgta |
| cPLA2 reverse | attcaccaccactaggcaGTCATGAAAGCGCTTTATGAATtacatctgtggcttcact |

Primers for RT-qPCR

| Name | Sequence |
|---|---|
| Chicken β-actin forward | ACAGCCCTGGCACCTAGCACAA |
| Chicken β-actin reverse | AAGGTGGACAGGGAGGCCAGGATA |
| Chicken PLA2g4a forward | CCCTCAATCTGACCAACAAGA |
| Chicken PLA2g5a reverse | CGGTAGATGAGGTCAAAGAG |
